# Pharmacodynamic model of PARP1 inhibition and global sensitivity analyses can lead to cancer biomarker discovery

**DOI:** 10.1101/2023.02.08.527527

**Authors:** Susan D. Mertins, Natalie M. Isenberg, Kristofer-Roy Reyes, Byung-Jun Yoon, Nathan Urban, Manasi P. Jogalekar, Morgan E. Diolaiti, M. Ryan Weil, Eric A. Stahlberg

## Abstract

Pharmacodynamic models provide inroads to understanding key mechanisms of action and may significantly improve patient outcomes in cancer with improved ability to determine therapeutic benefit. Additionally, these models may also lead to insights into potential biomarkers that can be utilized for prediction in prognosis and therapeutic decisions. As an example of this potential, here we present an advanced computational Ordinary Differential Equation (ODE) model of PARP1 signalling and downstream effects due to its inhibition. The model has been validated experimentally and further evaluated through a global sensitivity analysis. The sensitivity analysis uncovered two model parameters related to protein synthesis and degradation rates that were also found to contribute the most variability to the therapeutic prediction. Because this variability may define cancer patient subpopulations, we interrogated genomic, transcriptomic, and clinical databases, to uncover a biomarker that may correspond to patient outcomes in the model. In particular, GSPT2, a GTPase with translation function, was discovered and if mutations serve to alter catalytic activity, its presence may explain the variability in the model’s parameters. This work offers an analysis of ODE models, inclusive of model development, sensitivity analysis, and ensuing experimental data analysis, and demonstrates the utility of this methodology in uncovering biomarkers in cancer.

**Author summary:** Because biochemical reaction networks are complex, dynamic, and typically provide output that results from non-linear interactions, mathematical models of such offer insight into cell function. In the clinic, models including drug action further their usefulness in that they may predict therapeutic outcome and other useful markers such as those for prognosis. In this study, we report a model of drug action that targets a critical protein, that when inhibited, promotes tumor cell death and documented remissions. Because all patients do not respond to the described treatment, a means to find cancer patient subpopulations that might benefit continues to be a challenge. Therefore, we analyzed the pharmacodynamic model by defining the parameters of the greatest variability and interrogated genomic, transcriptomic, and clinical cohort databases with this information and discovered a novel biomarker associated with prognosis in some ovarian and uterine cancer patients and separately, associated with the potential to respond to treatment.

## Introduction

Cancer is a genetic disease manifest by both intra- and inter-tumour heterogeneity [1]. Further, clonal selection limits positive outcomes for patients as tumors, in many instances, generally do not show complete response to single therapeutic agent drugs and cells remaining after treatment may drive drug resistance. Signalling through biochemical networks, or signal transduction pathways, is complex and highly disrupted as well and can further contribute to tumor heterogeneity. Computational models can address these challenges as both normal and disease state can be captured in a fashion that closely resembles the dynamic *in vivo* milieu.

Pharmacodynamic modelling offers a means to assist with drug discovery and development. Modelling of reaction networks with ordinary differential equations (ODEs) has been an ongoing effort for decades [2] and has supplied insight into important principles of systems biology such as bistability, threshold responses, and the roles of positive and negative feedback loops [3–5]. Unlike AI models which have grown in popularity over the last few years, ODE models are tuneable and fully explainable, which offer a significant advantage. Oncogenic signalling has been modelled as well and has focused on cell fate outcomes such as cell cycle arrest and apoptosis [6,7]. In addition, models have been developed that address drug timing [8] and drug combinations [9].

Another specific use of pharmacodynamic models is the discovery of biomarkers that influence drug response in subpopulations of cancer patients, or in other words, aid in predictive oncology [10–12]. Hypothesis-driven experiments led to the discovery that cancer cells that have lost BRCA1/2 function for DNA repair are exquisitely sensitive to PARP1/2 inhibitors because PARP1 is critical in initiating the repair process. This synthetic lethality has been exploited clinically, confirming the biomarkers, BRCA1/2 [13]. However, not all patients with BRCA1/2 mutations have positive therapeutic outcomes and therefore, there is a need to define alternative biomarkers for sensitivity to PARP1 inhibitor treatment and/or prognosis in general. We present here a means to do so.

PARP1 is an enzyme that post-translationally modifies itself and target proteins by covalently adding chains of ADP-ribose units to specific amino acids [14,15]. It is activated by DNA strand breaks and, when activated, binds to sites of DNA damage where it triggers the assembly of DNA repair complexes. In normal cells, DNA damage occurs during DNA synthesis and transcription and PARP1 plays a key role in ensuring genomic integrity via repair mechanisms. In cancer, genomic instability can drive an increase in DNA breaks and, hence, a great dependence on DNA repair pathways. Within this context, inhibition of PARP1 can have therapeutic benefit. Here we describe an ODE model of PARP1 action, its inhibition, and a therapeutic output such as apoptosis abstracted to caspase activation. Subsequent sensitivity analyses identified and provided support for the role of a novel biomarker in RNA metabolism as it relates to translation. Extant databases containing clinical data were interrogated to determine if this same biomarker had clinical relevance and we uncovered it as a biomarker for prognosis. In summary, the computational methodology presented here led to the discovery of a previously unknown prognostic biomarker in RNA metabolism in ovarian and uterine cancer patients. Future development of the ODE model presented here and laboratory experiments may help define its clinical utility.

## Materials and methods

### Model development

Rule-based modelling represents cellular reaction networks in a dynamic fashion, encoding molecular interactions and associated kinetic rate parameters, while also accounting for site specific action such as phosphorylation at particular amino acids and other post-translational events [16,17]. The computational model reported here was written in the BioNetGen language, executed in an integrated design environment (RuleBender) which translates the reaction rules to ODEs, and thus, allows for large networks that are reliably created based on the defined parameters and interactions. Once defined, the ODEs can be solved either deterministically or stochastically for time-series outputs for species abundances contained within. The model is saved in a “bngl” format file (See S1 Text.) and provides a human-readable set of reaction equations for the process allowing those with little mathematical training easy access to understanding the biological basis of such models.

Several computational models have been put forward to describe cell fate decisions meditated by p53, namely cell cycle arrest or programmed cell death in the presence of genotoxic stress but did not describe a role for PARP1 initiated repair following DNA damage [6,7,18]. For the purposes of this effort, the ODEs reported by Bogdal *et al*. (referred to the Bogdal model herein) were coded using rule-based modelling and then adapted for PARP1 function and its binding to DNA double strand breaks [19]. The Bogdal model is important because it describes two well-described pathways leading to apoptosis as measured by caspase activation. The first is mediated by p53 abundance and its transcriptional activation of BAX, which, through a series of steps not fully included in the model, triggers caspase activation, the output. Baseline BAX is sequestered by binding to BCLXL. The second pathway is mediated by presumptive growth signals that, in turn, activate AKT, a kinase, which then phosphorylates BAD, keeping it bound to a scaffold protein and preventing its action in promoting programmed cell death. In the absence of AKT function, BAD is released and binds to BCLXL due to its higher binding affinity and displaces BAX from BCLXL, wherein BAX now ultimately triggers cell death. Thus. the absence of functional AKT results in caspase activation. Importantly, the authors demonstrate that the p53 and AKT pathways can act in a Boolean fashion dependent on the molecular abundances of BAD and BCLXL, where BCLXL can act as a buffer with tuning capabilities executing either ‘AND’ and ‘OR’ logic gates. In sum, depending on the molecular abundances of BAD and BCLXL, either or both pathways are required to trigger caspase activation. The newly adapted model presented in this report was coded as the ‘AND’ logic gate. Hence, the computational model was entitled the PARPi (PARP inhibitor) Logic Gate model.

Fig 1 is an extended contact map of the model reaction network that conforms to guidelines accommodating rule-based models without sacrificing critical details [20]. There are 14 molecules arranged for their location in the nucleus (white background) and the cytoplasm (shaded background) and 19 reaction rules numbered. The contact map describes the complete Bogdal reaction network plus additions, thus forming a new model not previously reported. DNA double strand breaks bind to either PARP1 or p53. It is well known that p53 is a transcription factor that responds to multiple genotoxic and other stresses, and in the case of the model, can trigger caspase activation through BAX mRNA synthesis. Therefore, competition between PARP1 and p53 is a simplification that supports our goal, to model the effect of PARP1 inhibitors, describe a therapeutic output, and provide a framework model for biomarker discovery. Further PARP1 function is also mapped by the addition of XRCC1, which is post-translationally modified by activated PARP1 and subsequently acted upon by PARG, an enzyme that removes poly-ADP (PAR) chains.

**Fig 1.**
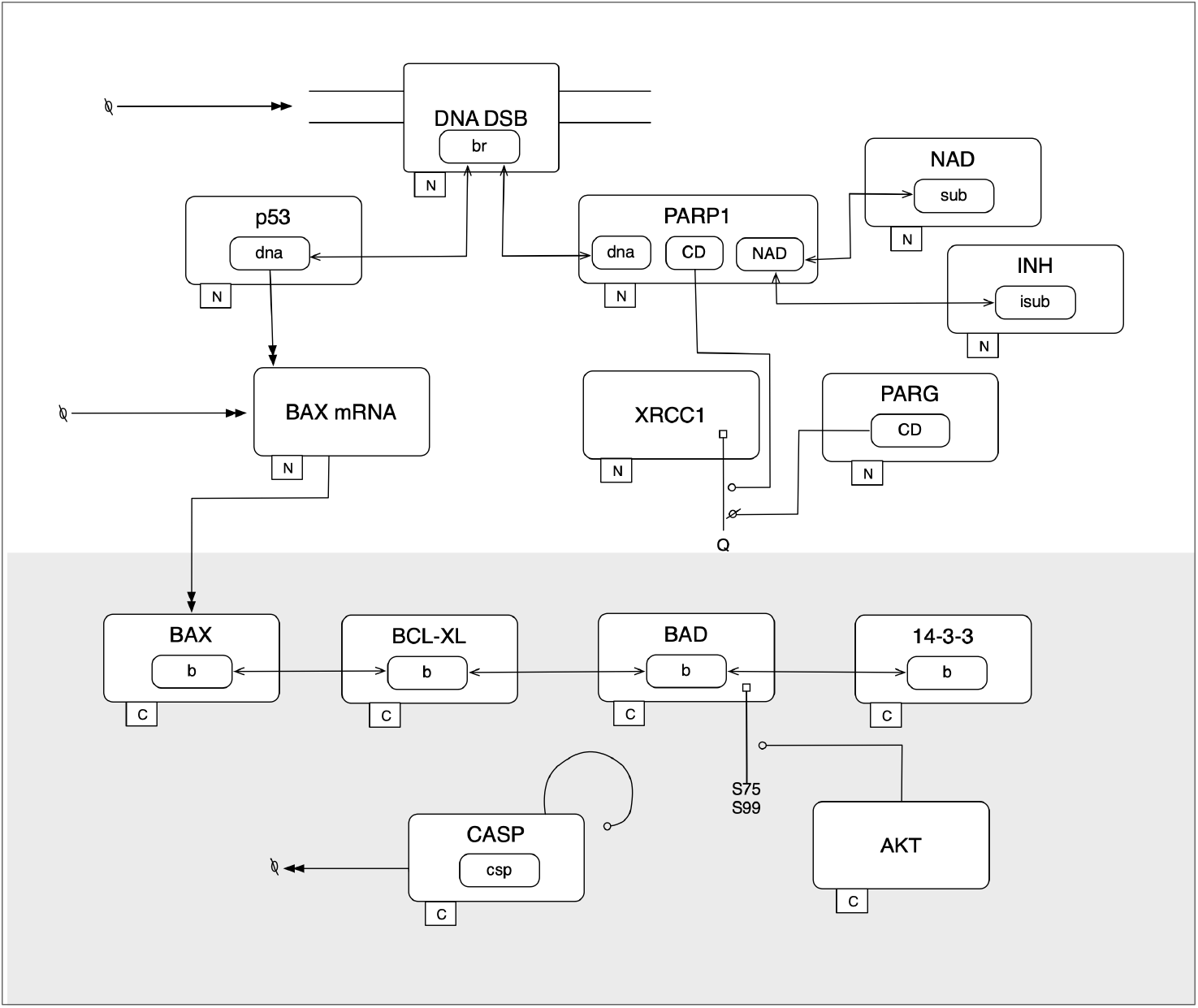
Extended contact map of PARPi Logic Gate Model. The Extended Contact Map is drawn from the model reaction rules (numbered, see S1 Table for specific code for the rules) following published guidelines [20]. Molecules are represented by rectangles containing components which may change state and interact in rules. Single-headed arrows (at both ends) depict binding reactions. Double-headed arrows describe synthetic reactions. Lines ending in open circles represent modifications. Flags originated in the molecules describe sites of modifications.

A complete description of the 19 reaction rules in the BioNetGen language can be found in S1 Table. Nearly all initial molecule concentrations and parameters (rate constants) were repeated from the report by Bogdal et al. [19] with the exception of DNA double strand breaks, PARP1, p53, PARG, NAD, and a generic inhibitor. These values were estimated and adjusted from reports measuring protein expression in cancer cell lines [21] and the metabolite, NAD [22,23]. S2 and S3 Tables supply all initial species concentrations and remaining parameters such as reaction rates, respectively. The complete rule-based model (bngl file in text format) can be found in S1 Text and is readily copied into RuleBender. Instructions for executing rule-based models and an updated IDE can be found at bionetgen.org. The model was simulated with a deterministic solver supplied by the IDE, RuleBender v2.3.0 (CVODE). From the 19 reaction rules, 30 ODEs were generated by BioNetGen that also elaborated 23 species. Runtime on a desktop computer (MacBook Pro, macOS Monterey 12.6.1, 8-core CPU, 16 GB memory), was minimal (CPU Time: 0.11s).

### Validation assay

Wild type and p53 mutant HCT116 cells (a gift from Fred Bunz [24]) were grown in McCoy’s 5A medium (Gibco, #16600-082) containing 10% fetal bovine serum (Corning, 35-010-CV) and 1% Penicillin and Streptomycin (Gibco, #15140-122) at 37°C and 5% CO_2_. Cell line identity was authenticated by short tandem repeat (STR) profiling and tested for Mycoplasma at Genetica. Short-term dose response assays were performed on the Incucyte^®^ ZOOM Live-Cell Analysis System (https://www.sartorius.com/en/products/live-cell-imaging-analysis/live-cell-analysis-instruments). To monitor cell proliferation and caspase activation, we generated Nuclight-labeled cells (Sartorius, #4625) and seeded cells in clear flat bottom tissue culture treated 384-well plate (Corning #3764) at a density of 200 cells/well (Volume: 50 μl/well). The next day, cells were treated with veliparib, olaparib, talazoparib at the indicated concentrations. 2-fold serial dilutions of each PARPi and etoposide were prepared in media containing the caspase-3/7 green dye (1:1000) (Sartorius, #4440) at starting concentrations of 10 μM, 10 μM, 2 μM, and 10 μM, respectively. Following treatment, plates were imaged every 4 hours over a period of 5 days and quantified using the IncuCyte^®^ ZOOM™ software algorithms to calculate phase confluence, red object count (reflecting cell count) and green object count (reflecting caspase activation). Fold change caspase activation was calculated as the ratio of the fraction of drug treated cells that were caspase positive compared to the fraction of DMSO-treated cells that were caspase positive. For dose response curves, all data were normalized to DMSO control. Non-linear regression and curve-fitting were performed in GraphPad Prism (Version 9.3.1).

### Sensitivity analysis

In order to evaluate the variance-based sensitivity of parameters in the model, the Sobol method, a global sensitivity approach, was conducted [25]. The Sobol method assumes that the variance in model output can be attributed to subsets of inputs, from all single parameters to all the pair-wise contributions, and so on.

As indicated above, the model was represented by a system of coupled, non-linear ODEs containing 27 parameters (S3 Table). We provide lower and upper bounds in S4 Table which represent 90% confidence intervals for each of the parameters within the model. These intervals indicate our expert knowledge regarding the lower and upper limits indicating where 90% of the probability mass lies for each uncertain parameter. Further details regarding the origins of these limits can be found in S2 Text. The model and global sensitivity analyses were implemented in Julia [26] using the DifferentialEquations.jl and GlobalSensitivity.jl libraries, making use of powerful ODE solvers and diagnostics [27,28]. For the Sobol sensitivity analysis, IC50 was fixed to a value of 0.01 and was not considered uncertain. For tractability, 1,000 samples were used in computing the estimators in all Sobol indices.

To utilize a variance-based sensitivity analysis such as the Sobol method, a quantity of interest (QoI) was defined. QoI is a metric computed from the output of the ODE model that will be used to quantify the impact of parameter uncertainty on the model. For this study, we select maximum active caspase concentration as the QoI, where maximum active caspase concentration is defined as 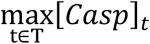, where *t* ∈ *T* are the set of time points for integrating the ODE, and [*Cαsp*]_*t*_ is the active caspase concentration at time *t*. The maximum activated caspase concentration achieved over the time course of the simulation of the ODE is expected to relate to the model prediction of cell apoptosis.

### Bioinformatic analysis for biomarker discovery

Several well-established and curated databases were interrogated to uncover possible biomarkers consistent with the findings of the global sensitivity analysis. First, the Gene Expression Omnibus (https://www.ncbi.nlm.nih.gov/geo) was searched for previously published studies that quantified transcriptomics of cancer samples (*in vivo* and *in vitro*) following treatment with PARP1 inhibitors [29]. The work by Sheta *et al*. was selected which provided a list of differentially expressed genes in cultured ovarian tumor spheroids sensitive to PARP1 inhibition (see S5 Table for the list.) [30].

Gene set enrichment was conducted at two separate databases, GSEA (https://www.gsea-msigdb.org/gsea) [31,32] and GO Resource (http://geneontology.org) [33,34] to determine if there was a preponderance of genes that associate with protein synthesis and/or RNA metabolism. Statistically significant groupings were captured. Inspection of the individual genes in particular groups for functions consistent with parameters uncovered in the global sensitivity analysis revealed a gene, GSPT2 with these characteristics.

GSPT2 (HUGO ID: HGNC:4622) was subsequently studied at the cBioPortal for Cancer Genomics database (https://www.cbioportal.org) [35–37]. cBioPortal database is a large and curated compendium of cancer patient cohorts reported in the literature that readily allows study of a gene of interest or cancer type. Queries for genomic and transcriptomic features and selected subpopulations were conducted. Data included copy number variation, mRNA expression levels, mutational events (including exact sites and amino acid changes), and overall survival. These data types were collected from ovarian cancer patients (TCGA, Firehose Legacy) and uterine cancer patients (TCGA, PanCancer Atlas).

Statistical tests for analyses in the cBioPortal database were conducted and included those for overall survival (Wilcoxon signed-rank test), mutation frequency (X^2^ test), and differences in mRNA expression levels (*t* test) where a = 0.05.

The PRECOG database catalogues and combines transcriptomic profiles from cancer cohorts and overall survival for analysis (PREdiction of Clinical Outcomes from Genomic Profiles (PRECOG, https://precog.stanford.edu/) [38]. GSPT2 expression was queried to obtain a Z-score through a meta-analysis across many cancer histologies and within a particular histology for a confirmatory analysis.

## Results

### Model simulation

The Boolean logic gate model reported by Bogdal *et al*. initiated apoptotic signalling via two well-described induction pathways, p53 and inactive AKT. In the report, the authors demonstrated that, dependent on initial concentrations of certain species, either one pathway or both pathways can cause caspase activation. Model output was coded through BAX upregulation and caspase activation as a representation of programmed cell death. Initial steady state concentrations (in molecule number or abundances) of p53 and AKT were manipulated in a multistep analysis in order to demonstrate that altering molecular abundances of BAX can bring about the need for either (‘OR’ gate) pathway or both (‘AND’ gate) pathways to trigger caspase activation.

Specifically, the Bogdal model demonstrated the OR gate where either inactive AKT or p53 levels representing a response to DNA damage trigger caspase activation. Further, altered molecular abundances of BAD (approximately 2.5-fold lower than those utilized in the OR gate model) required both inactive AKT and p53 to trigger caspase activation. This latter model was designated an ‘AND’ gate. For the purposes reported here, when the exact ODEs (in the ‘AND’ gate, reported in the Bogdal model) were translated to a rule-based model, identical output was found (S1 Fig). Of note, these simulations were run prior to the changes described below.

As a first step, the Bogdal model was amended to include PARP1 function and its inhibition. This simple model reported here was expected to provide background for subsequent analyses leading to biomarker discovery. It should also be noted that our model can be advanced to include important phenomena such as synthetic lethality under homologous recombination deficiency in cancer. The following changes were included: 1) DNA double strand breaks as a species were added for competitive binding between p53 and PARP1.; 2) PARP1 bound its substrate NAD+ and PARylated a representative protein, XRCC1.; 3) PARG, a poly ADP-ribose glycohydrolase, was included to balance PARP1 signalling.; 4) A species for a PARP1 inhibitor was included that competed with NAD+ binding.; 5) IC50 values for the inhibitor were considered as a rate constant when multiplied by the dissociation constant for NAD+ binding to PARP1. Each IC50 value was derived from *in vitro* functional assays and represents all possible inhibitors, known and unknown. (See Fig 1 for reaction network overview (or extended contact map) and S1-S3 Tables for code defining the rules and associated parameters.)

Once the PARPi Logic Gate model was amended to include the effects of an inhibitor and its competitive binding with NAD+, simulations were run to determine if caspase activation occurred, consistent with a rise in BAX molecular abundance (or molecule number). It should be noted that, in the Bogdal model, when BAX molecule abundance exceeded 5000 molecules over 10 hours, caspase was activated in an ultrasensitive response. In addition, activated caspase abundance above 1000 molecules was considered sufficient for apoptotic induction. Taken together, in our model, Fig 2A (BAX abundance) and Fig 2B (activated caspase abundance) demonstrated a dose response to a PARP1 inhibitor (IC50 ranges scanned were from 0.0001 – 10.0 μM). Activated caspase abundance occurred in the range of 0.0001 – 0.01 μM. Importantly, when the inhibitor was removed from the model, simulation output shown in Fig 2C demonstrated that BAX levels were not sufficient to trigger caspase activation (BAX abundance is approximately 250 molecules). Fig 2D provides further support that the reaction network is consistent with the output goals. Notably, in the absence of an inhibitor, XRCC1 was found to be in the PARylated state and this condition was greater than when PARP1 was inhibited by 0.001 μM coded as the IC50 concentration.

**Fig 2.**
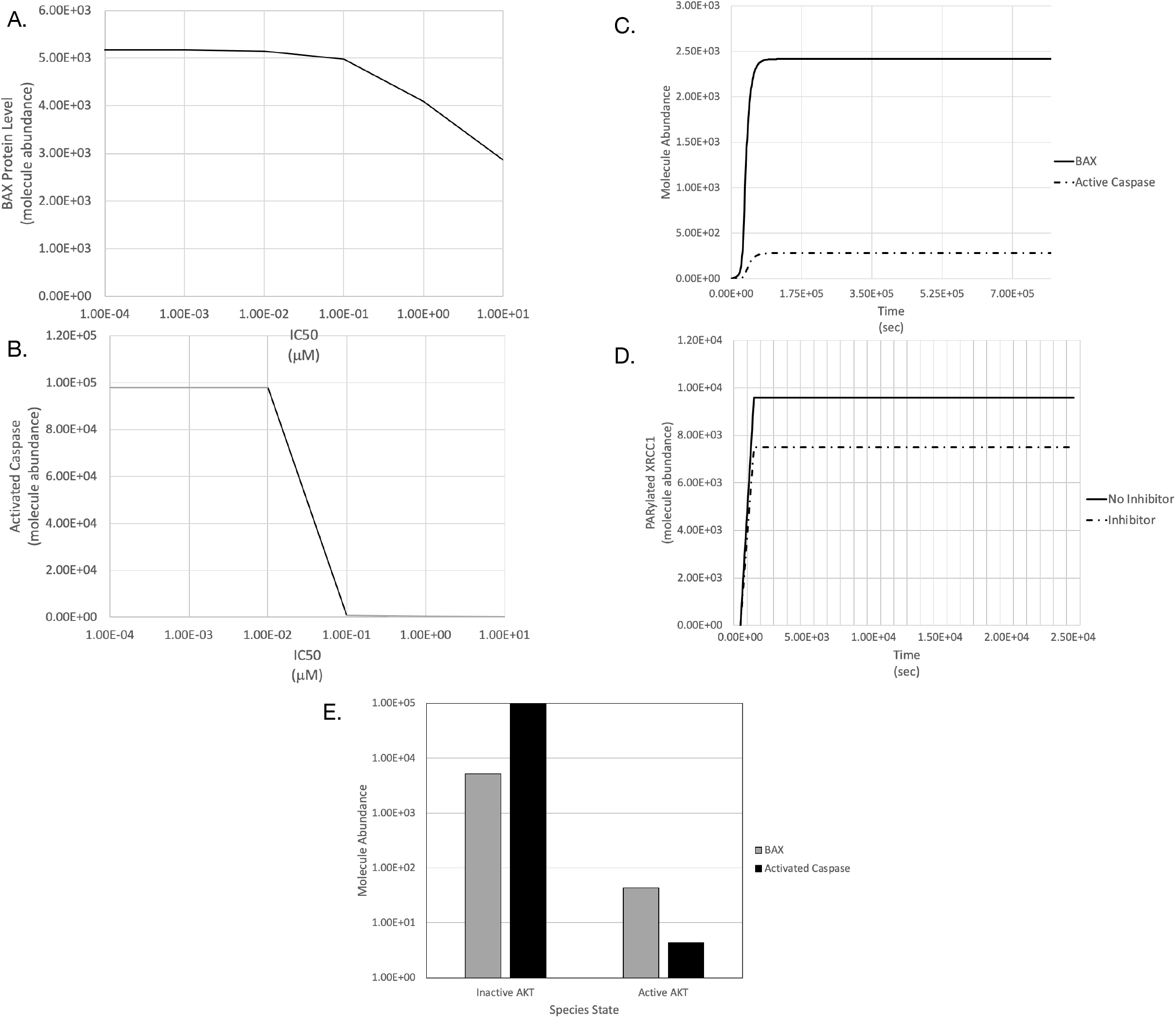
Model simulation output. Simulations were run using the deterministic CVODE provided with the BioNetGen IDE, RuleBender. A. The level of BAX protein is reported for the range of IC50 concentrations evaluated in the model. As previously reported, 5000 or more molecules of BAX will trigger caspase activation [19]. B. Activated caspase occurs at low doses of a generic inhibitor. C. When no inhibitor is present in the model, BAX protein molecule abundance does not exceed 5000 and caspase is activated to background levels. D. XRCC1, a substrate for PARP1 PARylation occurs when no inhibitor is present and is reduced in the presence of an inhibitor at 0.001 μM. E. To demonstrate that caspase activation is dependent on the p53 pathway in the model (OR gate), both inactive and active AKT was evaluated in the model. With inactive AKT (i.e., no phosphorylated AKT), BAX and activated caspase are induced. With active AKT, nearly no BAX or activated caspase was formed. Thus, the model acts as previously published.

Because the PARPi Logic Gate model presented here is an advance on the ‘AND’ logic gate model presented by Bogdal *et al*., it was also important to confirm that both sufficient p53 levels and inactive (unphosphorylated) AKT were required for caspase activation. In the model as reported in S1 Text, unphosphorylated AKT is represented by an initial concentration of zero. Therefore, an increase in initial concentration for AKT represents a level of activated (phosphorylated) AKT and thus, was expected to reduce or inhibit caspase activation. Fig 2E supports this expectation. When 2 x 10^4^ AKT molecules were added to a simulation run, BAX abundance was insufficient to trigger an increase in caspase activation (i.e., below 1000 BAX molecules). Because it has been reported that mutant p53 typically accompanies effective PARPi therapy in BRCA1 mutant breast cancer, future studies will address this genotype [39].

### Validation data

To provide support for the PARPi Logic Gate model and its therapeutic output, we performed *in vitro* growth inhibition studies and measured caspase activation in HCT116 colon tumor cells exposed to 3 clinically approved PARPi with varying IC50 values, talazoparib, olaparib, and veliparib (Fig 3 and S3 Fig). Several lines of evidence aided in validating the model. First, we noted an increase in caspase3/7 levels in cells treated with PARPi, indicating that the well-defined apoptotic cell death pathway can operate under PARP1 inhibition. Secondly, caspase activation (approximately 2-fold) occurred at day 3.8 at the IC50 concentration in talazoparib and olaparib treated cells (Fig 3). This is consistent with the ODE model where caspase activation occurred at approximately day 3.5 at IC50 values equal to or below 0.03 μM. Thus, the timeline of caspase activation was similar between our model and experimental data. Third, the experimental data demonstrated that caspase activation correlated with potency with more potent agents (i.e., talazoparib) showing caspase activation at lower doses (Fig 3). Thus, the model sigmoid dose response curve in Fig 2B for a PARP1 inhibitor was consistent with the shape of the dose response curves reported for the 3 experimentally evaluated drugs (talazoparib – 0.0042 μM, olaparib – 1.25 μM, and veliparib – 15.19 μM) (S3 Fig). Finally, it was notable that the ODE model presented here was an adaption of one widely investigated previously and validated in other respects [7,19,40].

**Fig 3.**
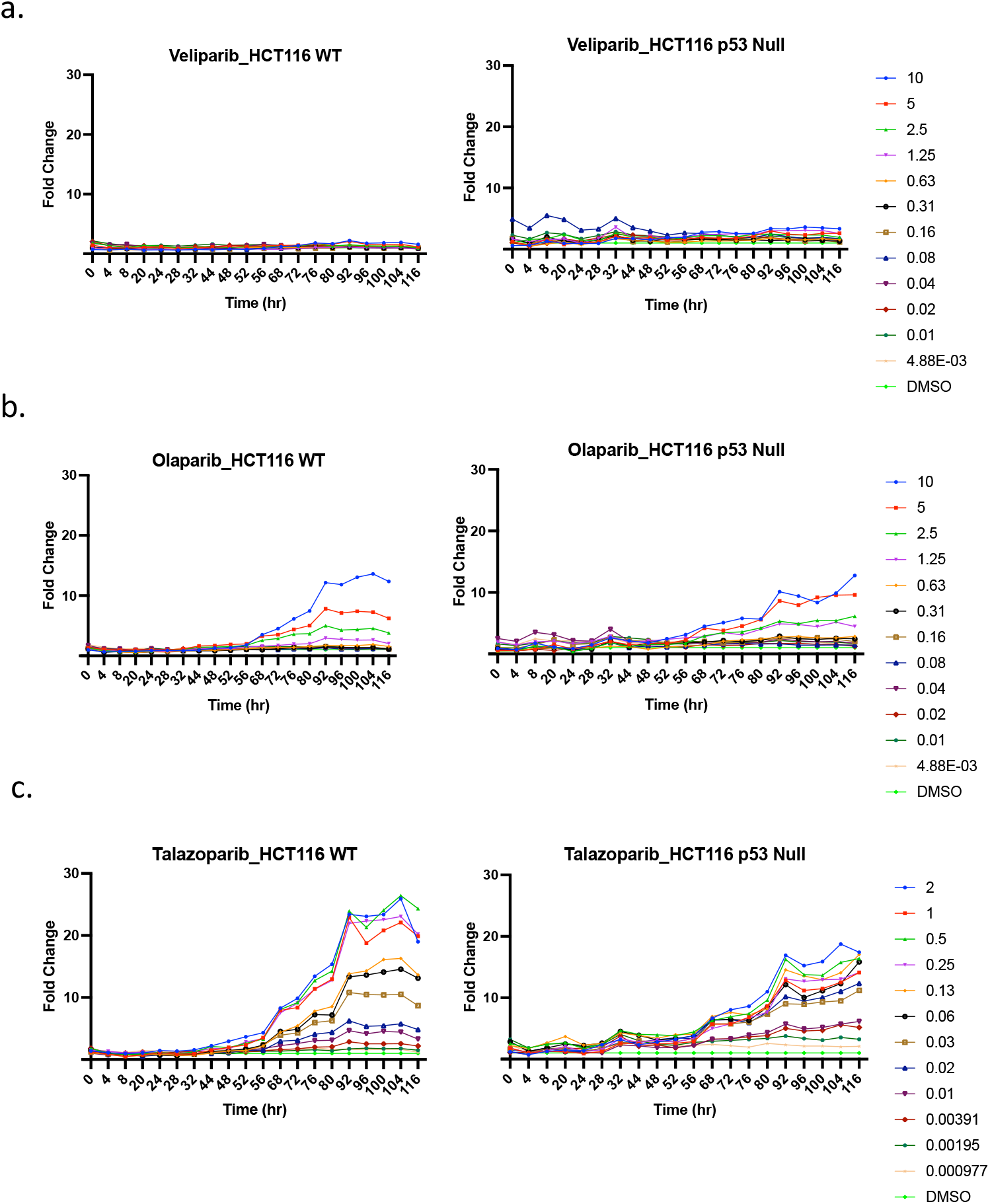
Effect of three clinical PARP1 inhibitors on caspase activation in HCT116 cells. The relative fraction of caspase positive HCT116 WT (left) and p53-/- (right) cells over time after being treated with the indicated concentrations of (a) veliparib, (b) olaparib, and (c) talazoparib. Caspase activation was calculated as the ratio of the fraction of drug-treated cells that were caspase positive compared to the fraction of DMSO-treated cells that were caspase positive at each time point.

### Sensitivity analysis findings

Results for the first-order and total-effect Sobol indices are summarized in Fig 4 for each parameter utilizing the QoI of maximum activated caspase. The first-order Sobol index represents the effect on the variance of each individual parameter on the variance in the model QoI, averaged over the variations in all other parameters. The total-effect represents the contribution to the variance of the QoI imparted by each individual parameter alone, as well as its contributions due to all higher-order interactions between other parameters. Together, the first-order and total-effect Sobol indices can give insight into (1) which uncertain model parameters and (2) which uncertain model parameter interactions, contribute the most to the overall uncertainty in the model output. This, in turn, gives insight into which model parameters warrant further experimental study to reduce uncertainty regarding their values.

**Fig 4.**
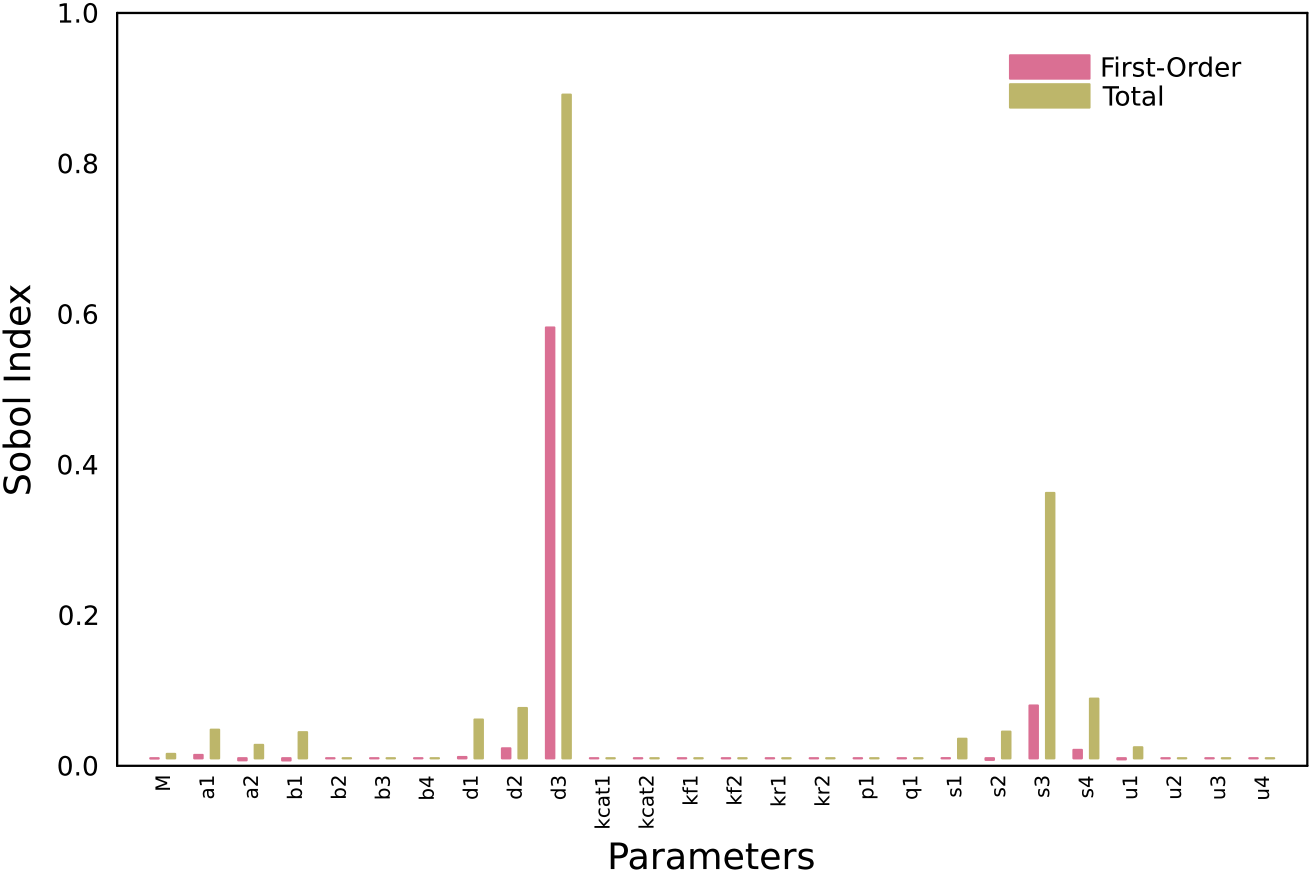
First-order and total-effect Sobol index values for each parameter in the PARPi Logic Gate Model. Higher first-order and total-effect Sobol index values indicate larger impact of the parameter variance (independently and over all parameter interactions, respectively) on the variance in the model output.

As seen in Fig 4, the largest Sobol index values were seen in parameters *d*_3_ and s_3_. More interestingly, both parameters have higher total-effect indices when compared to their first-order indices. This implies that there were some higher order interactions in these two parameters which must be contributing to the variance in the QoI of the model.

To confirm this, the second-order Sobol indices were computed, which is defined as the contribution of all pairwise parameter interactions to the model output variance. The computation of the second-order Sobol index generates 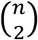 results, where *n* is the number of model parameters, and very few of the computed second-order indices have large enough magnitudes to be regarded as significant contributions. However, it was notable that the second-order Sobol index for the pairwise contribution of *d*_3_ and *s*_3_ was two orders of magnitude greater than any other pairwise interaction in the model with a value of 0.22. This implied that the QoI responds nonlinearly when the two parameters were varied together, and that reducing the uncertainty regarding the values of *d*_3_ and *s*_3_ would greatly improve the confidence in predictions made by the model.

### Bioinformatic analysis

Because the sensitivity analysis demonstrated that two parameters, s3 (*s_3_*) for pro-caspase synthesis rate and d3 (*d_3_*) describing pro-caspase and caspase protein degradation rate contributed to the most variation in the model, it may indicate that there are biomarkers for RNA metabolism that determine PARP1 inhibitor sensitivity and/or resistance. Recently, a role for nonsense mRNA mediated decay in cancer has been delineated and, therefore, provided a rationale for this hypothesis [41]. Further, it would be expected that within this variation, one can find patient subpopulations better suited for treatment with such clinical agents. In addition, an enzyme that manifests its activity through changes in reaction rates or binding affinity with substrates or molecular partners would be of interest and support a dynamic model as presented here.

A previously published report included a list of genes over- and under-expressed in ovarian tumor spheroids treated with the PARP1 inhibitor, olaparib [30]. Altered expression of 602 mRNA expression levels were associated with a PARP1 inhibitor sensitive phenotype and are listed in S5 Table. Two separate enrichment analyses for biologic pathways were conducted. In the first study, limited by the algorithm, the top 500 over- and under-expressed genes were examined for pathway enrichment using GSEA (Gene Set Enrichment Analysis, https://www.gsea-msigdb.org). The pathways considered for enrichment were defined by Reactome, KEGG, BioCarta, and Wikipedia Pathways [42–45]. Significant pathway associations are listed in Table 1. While none of the significant pathways found specifically addressed protein synthesis and RNA metabolism, the genes within the immune system and nervous system development categories were examined individually for function in RNA metabolism as the process may be critical in immune regulation and nervous system development. For instance, viral mRNA degradation is a step that requires cellular machinery for such action and separately, neurons have challenges over intracellular distances in translational machinery, in particular, regulating mRNA stability from point of production to site of need. The genes enriched in nervous system development were individually examined and are listed in Table 2. GSPT2 (G1 to S Phase Transition 2), also known as ERF3b (Elongation Release Factor 3b) (HUGO ID HGNC:4622), was considered for further study due to its potential role in RNA metabolism and regulation of cell cycle progression. GSPT2 is a GTPase that functions in both promoting efficient ribosome initiation and elongation termination [46]. Therefore, considering the sensitivity analysis findings, GSPT2 may influence both protein synthesis and protein degradation rates as well as the licensing of cell cycling. GSPT2 was overexpressed (2.18-fold) in olaparib sensitive ovarian tumor spheroids [30]. In the Sheta report, GSPT2 was ranked the 92^nd^ most upregulated gene (of 239 such genes) with greater than 1.5-fold increase (p < 0.05). A complete list of significant over- and under-expressed genes can be found in S5 Table. Inspection of the gene list in the GSEA analysis with immune regulation function did not reveal any genes with a role in RNA metabolism.

**Table 1.** Significant Pathway Association using the Gene Set Enrichment Analysis (gsea.broad.org).

| Gene Set Name | Pathway Source | p-value | FDR q-value |
| --- | --- | --- | --- |
| Matrisome | Naba et al. | 1.25E-10 | 2.03E-07 |
| Developmental Biology | Reactome | 1.36E-10 | 2.03E-07 |
| Extracellular Matrix Organization | Reactome | 4.46E-10 | 4.44E-07 |
| RHO GTPase Cycle | Reactome | 8.06E-09 | 5.06E-06 |
| Matrisome Associated | Naba et al. | 8.49E-09 | 5.06E-06 |
| Nervous System Development | Reactome | 2.49E-08 | 1.24E-05 |
| Signaling by RHO GTPases, Miro GTPases, and RHOBTB3 | Reactome | 6.34E-08 | 2.70E-05 |
| Glucocorticoid Receptor Pathway | WikiPathways | 2.81E-07 | 1.05E-04 |
| Hemostasis | Reactome | 3.65E-07 | 1.15E-04 |
| Signaling by Receptor Tyrosine Kinases | Reactome | 3.86E-07 | 1.15E-04 |
| Cell Adhesion Molecules | KEGG | 4.30E-07 | 1.17E-04 |
| Malignant Pleural Mesothelioma | WikiPathways | 9.16E-07 | 2.28E-04 |
| Post-translation Protein Modification | Reactome | 1.28E-06 | 2.94E-04 |
| Cell Surface Interactions at the Vascular Wall | Reactome | 1.90E-06 | 3.92E-04 |
| Infectious Disease | Reactome | 1.97E-06 | 3.92E-04 |
| Prostaglandin Synthesis and Regulation | WikiPathways | 3.28E-06 | 5.78E-04 |
| VEGFA - VEGFR2 Pathway Signaling | WikiPathways | 3.30E-06 | 5.78E-04 |
| Cytokine Signaling in the Immune System | Reactome | 3.51E-06 | 5.82E-04 |
| A6B1 and A6B4 Integrin Signaling | Pathway Interaction Database | 3.75E-06 | 5.88E-04 |
| MHC Pathway | BIOCARTA | 6.27E-06 | 9.35E-04 |

**Table 2.** Genes Enriched in Nervous System Development.

| Abbreviated Gene Name | Gene (HUGO ID) |
| --- | --- |
| ITGAV | integrin subunit alpha V [HGNC:6150] |
| ITGA1 | integrin subunit alpha 1 [HGNC:6134] |
| ABL2 | ABL proto-oncogene 2, non-receptor tyrosine kinase [HGNC:77] |
| MYL9 | myosin light chain 9 [HGNC:15754] |
| SHC1 | SHC adaptor protein 1 [HGNC:10840] |
| FGFR1 | fibroblast growth factor receptor 1 [HGNC:3688] |
| NRP1 | neuropilin 1 [HGNC:8004] |
| NEO1 | neogenin 1 [HGNC:7754] |
| PSMB8 | proteasome 20S subunit beta 8 [HGNC:9545] |
| PSMB9 | proteasome 20S subunit beta 9 [HGNC:9546] |
| ROBO1 | roundabout guidance receptor 1 [HGNC:10249] |
| RGMB | repulsive guidance molecule BMP co-receptor b [HGNC:26896] |
| MBP | myelin basic protein [HGNC:6925] |
| RPS6KA6 | ribosomal protein S6 kinase A6 [HGNC:10435] |
| CD24 | CD24 molecule [HGNC:1645] |
| DPYSL4 | dihydropyrimidinase like 4 [HGNC:3016] |
| **GSPT2** | **G1 to S phase transition 2 [HGNC:4622]** |
| PDLIM7 | PDZ and LIM domain 7 [HGNC:22958] |
| DOK6 | docking protein 6 [HGNC:28301] |
| LAMB1 | laminin subunit beta 1 [HGNC:6486] |
| NTN4 | netrin 4 [HGNC:13658] |
| SLIT2 | slit guidance ligand 2 [HGNC:11086] |

In the second enrichment analysis, the Gene Ontology Resource (http://www.geneontology.org) was queried with the complete list of 602 over- and under-expressed genes in ovarian tumor spheroids to determine if enrichment occurred for GO terms. Gene Ontology is a knowledge base that defines genes through selective and curated terminology that takes a parent and child relationship in a graph [33,34]. The terms are biologic process, molecular function, and cellular location. The results demonstrated that for 598 out of 602 genes, there was a significant enrichment across 380 biologic processes of varying parent-child relationships (Complete data set not shown but available on request). Overall, there were similarities to the GSEA findings reported above (for immune and nervous systems) except that mRNA metabolism genes were found to be significantly under-represented (for RNA splicing, p = 6.12e-4, False Discovery Rate (statistical correction for multiple comparisons, FDR) = 2.91e-2 and for mRNA processing, p = 5.51e-5, FDR = 4.83e-3).

To determine if there was clinical relevance for GSPT2 as a biomarker for cancer prognosis and/or treatment, the cBioPortal for Cancer Genomics database was interrogated (https://www.cbioportal.org) [35–37]. This open-access online resource includes genomic, transcriptomic, and clinical data from a large number of clinical cohorts, allowing for investigations into mutational status, overall survival, and subpopulation characterization. Specifically, queries by gene or explorations by selected clinical cohort can be analysed. The Cancer Genome Atlas (TCGA) cohort of 617 patients with ovarian cancer was studied. In this population, 23 patients possessed GSPT2 amplifications and had worsen overall survival (Table 3, Z = −2.48, p = 0.007). Another gene that showed amplification in ovarian tumors (NOCL2) with a similar frequency as GSPT2 was randomly chosen to determine if gene amplification, in general, could predict worsened overall survival. The Wilcoxon Signed Rank Test was conducted and no significant results were obtained (Z = 0.74, p = 0.23).

**Table 3.**
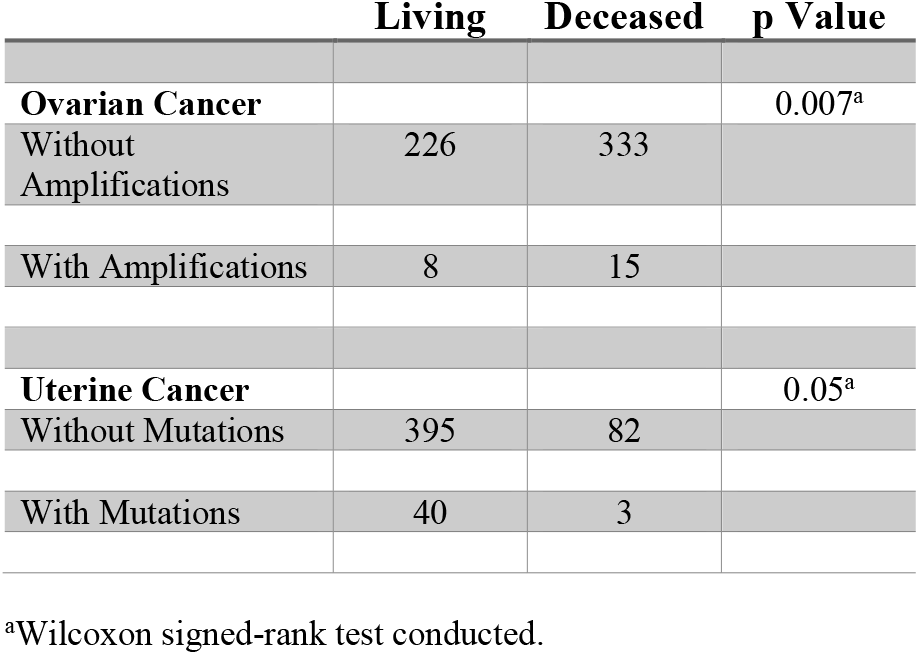
Overall Survival in Ovarian and Uterine Cancer Patients (TCGA Cohort) with GSPT2 Aberrations.

|  | **Living** | **Deceased** | **p Value** |
| --- | --- | --- | --- |
| **Ovarian Cancer** |  |  | 0.007<sup>a</sup> |
| Without Amplifications | 226 | 333 |  |
| With Amplifications | 8 | 15 |  |
| **Uterine Cancer** |  |  | 0.05<sup>a</sup> |
| Without Mutations | 395 | 82 |  |
| With Mutations | 40 | 3 |  |

<sup>a</sup>Wilcoxon signed-rank test conducted.

GSPT2 was not found to contain any mutational defects, however, we did note a significant decrease in GSPT2 mRNA expression compared to patients with the diploid complement of GSPT2 (p = 0.003, Fig 5) in the TCGA cohort, surprisingly. To confirm this result, the PRECOG database was queried for any association between GSPT2 expression and overall survival in ovarian cancer patients. Although statistical significance was not found, in this instance, 13 datasets were combined to support a trend that lower expression of GSPT2 was associated with poorer overall survival (Z = 1.29, p = 0.10). In conclusion, for ovarian cancer patients, there was support for GSPT2 over-expression as a biomarker for PARP1 inhibitor sensitivity [30] but separately, with regards to amplification, as a prognostic factor as observed in the clinical cohort. Future studies will be needed to confirm overall decreased GSPT2 gene expression in patients with GSPT2 amplification.

**Fig 5.**
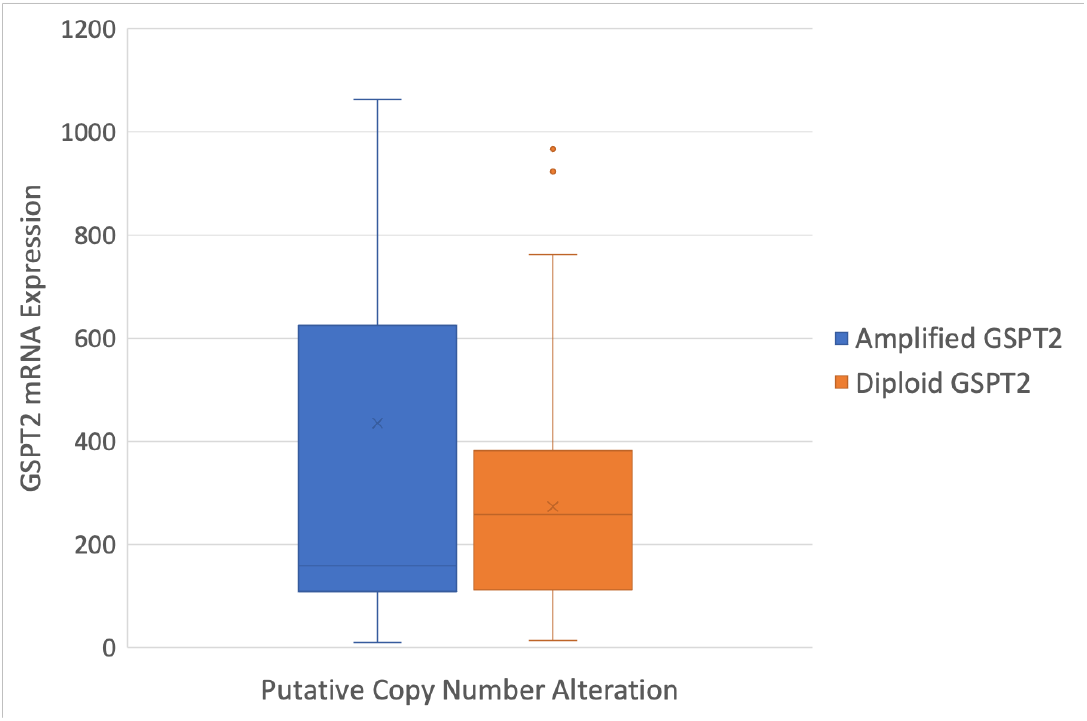
Putative copy number in ovarian cancer cohort (TCGA cohort) reported in cBioPortal.org. Patients with amplified tumor GSPT2 had a statistically significant decrease in GSPT2 mRNA compared to patients with a diploid copy number. Note the box and whiskers plot reporting 75^th^ and 25^th^ percentiles at the bars.

In an investigation of the uterine cancer cohort from the TCGA (Pan Cancer Atlas, 529 patients), 8.3% of patients had GSPT2 missense or truncating mutations. S1 Fig combines images taken from Pfam (https://pfam.xfam.org) describing the GSPT2 domain structure [47]. Further study of the particular mutations revealed 5/39 patients had mutations in the N-terminal domain (PAM2 domain (binding PABP scaffold protein), 25/39 had mutations in P loop (of the GTPase domain), and 9/39 had mutations in the C-terminal domain (critical for elongation termination function). A chi-squared test revealed that it was not likely that the distribution of missense mutations was due to random events (p < 0.005, df =2, X^2^ = 83.47). Furthermore, patients with mutations had improved overall survival (Table 3, Wilcoxon rank-sum test, Z = 1.65, p = 0.05). It is possible that selection for certain mutations in GSPT2 in uterine cancer patients can limit detrimental aspects of carcinogenesis and overcome the oncogenic processes. (N.B. One patient of 39 with mutations had both GSPT2 amplification and documented mutation (C93G).)

## Discussion

A mathematical model using rule-based code was developed that described PARP1 functional inhibition by a generic modifier and its therapeutic outcome as measured by caspase activation. This was an advance of a previously published model that defines cell fate dependent on two pathways capable of triggering programmed cell death in the presence of DNA damage and/or the absence of growth factors (p53 or AKT pathway, respectively). Qualitatively, the model was confirmed with laboratory experiments that describe dose responses to clinical agents and timing of the apoptotic outcome. A global sensitivity analysis pinpointed two rate constants describing protein synthesis and degradation as contributing to the most variation in caspase activation predictions. Finally, it was hypothesized that a gene or genes may be discovered through bioinformatic analyses that, functionally, may explain the newly discovered variable rate constants. GSPT2, a GTPase known to both promote ribosome initiation and terminate protein synthesis, was identified to be a prognostic biomarker, but with opposing effects on overall survival in ovarian and uterine cancer cohorts. New computational models that incorporate a role for GSPT2, its enzymatic activity, and binding partners, and future wet laboratory experiments will help determine the potential impact of the findings in this report.

The PARPi Logic Gate model presented here capitalizes on a previous report by Bogdal *et al*. [19]. That model was limited in scope but was useful for the present purposes, as it readily allowed the addition of PARP1 DNA binding action and its catalytic activity to be incorporated alongside p53, a critical transcription factor that triggers downstream outcomes in face of genotoxic stress such as DNA damage. It is notable that most published mathematical models describing p53 action focus on its induction after DNA damage via irradiation. Yet, presently, no mathematical models include the functional activity of PARP1. Therefore, our model is forming the baseline for future studies where significant details of downstream action of PARP1 are incorporated, mainly its role in repairing DNA double strand breaks via homologous recombination. Further, incorporating mutational status of this machinery such as BRCA1/2 aberrations may help build a model for synthetic lethality.

The mathematical model was validated qualitatively as a first effort, but importantly, the laboratory results confirm that the levels of caspase activation predicted by the PARPi Logic Gate model correlate with the range of IC50 values of clinical agents (veliparib IC50 > olaparib IC50 > talazoparib IC50) measured empirically. In addition, the induction of caspase activation (3.5 days) in the model approximates that found for the laboratory experiments (3.8 days). It will be of interest to consider intracellular drug concentration and other measurable output to support the model such as the formation of RAD51 foci due to action of the homologous recombination machinery following drug exposure.

Sensitivity analyses have been commonly used following construction of computational ODE models to discover potential drug targets, therapy resistance, and other biomarkers for clinical application (for example, see [48–51]). Here, two related rate constants that direct protein synthesis and degradation were found to contribute to the most variability in model output, caspase activation. Recently, Kardynska *et al*., presented a novel sensitivity analysis utilizing a p53 network model without explicit apoptotic reaction rules and discovered similar sensitive parameters affecting transcription and translation rates as critical drug targets to modulate p53 induction of programmed cell death [52]. Importantly, for this study, we focused on the parameters affecting caspase activation and the role of PARP1 inhibition and we further evaluated biomarkers of PARP1 inhibitor sensitivity or resistance and/or prognostic factors. This type of investigation (i.e., linking sensitivity analysis findings and bioinformatic database interrogations) has not been undertaken before and therefore, implicitly offers the incorporation of both dynamic behavior of cell signalling and the heterogeneity found in tumors in the subsequent outcomes.

It is well known that PARP1 inhibitors are synthetically lethal with loss of BRCA1/2 [13]. Further, this paradigm has evolved to encompass those molecules contributing, overall, to homologous recombination, so-called homologous recombination deficient phenotypes (HRD) [53]. Our model and global sensitivity analyses pointed to other biomarkers other than those relating to the HRD phenotype. Utilizing data from a published study, we identified GSPT2 as a presumptive biomarker of PARPi inhibitor sensitivity in ovarian tumors [30]. GSPT2 (or ERF3b) is a GTPase that has catalytic activities consistent with protein synthesis and degradation. Within a complex assembled at the ribosome, GSPT2 promotes protein synthesis by allowing efficient ribosome initiation. It has also been shown to modify mRNA elongation termination [46]. Therefore, the addition of GSPT2 function in the PARP1 Logic Gate model may improve its predictions and, dependent on its catalytic rate *in vivo*, explain PARP1 inhibitor sensitivity. In particular, slower elongation termination for caspase mRNA may increase sensitivity while faster or premature termination might remove caspase mRNA and limit programmed cell death induction.

Interrogating bioinformatic databases offered the opportunity to determine if GSPT2 had any clinical relevance. Therefore, we examined clinical data sourced from ovarian and uterine cancer cohorts. We found that GSPT2 amplification may worsen overall survival in patients with ovarian cancer but associated with improved survival in uterine cancer patients without amplification, but rather possessing mutated GSPT2. These contradictory findings may reflect the dual functions ascribed to GSPT2 noted above. Thus, the findings presented on these clinical cohorts are encouraging and support the other analyses, but warrant closer examination, for both as a role in overall survival and therapeutic outcome.

For uterine cancer patients, GSTP2 mutations appear to support overall survival, suggesting a mutational profile that dampens oncogenesis. Future studies may examine one of the recurrent mutations as one was found more frequently in the presumptive P loop of the GTP domain (E270K, in 4 of 39 patients). This location, near the region where GTP is predicted to enter the active site, is of interest and it bodes for a change in the rate constant. Therefore, molecular dynamic and docking studies may be of interest. How improved survival is mediated is unclear, but a better understanding for a dual role of mRNA decay, in both promoting and supressing oncogenesis is now emerging in cancer [41].

In summary, we report an advance that incorporates drug action on PARP1, a critical initiator of the DNA damage response on a previously published mathematical model of apoptosis signalling. In addition, we pinpointed a biomarker by exploiting findings from a global sensitivity analysis and interrogations of bioinformatic databases and we await future confirmatory laboratory experiments. In the future, the mathematical model presented in this paper may be integrated with generative artificial intelligence models for *de novo* molecular design [54–56] of an enhanced PARP1 inhibitor with desirable therapeutic outcome such as caspase activation.

## Supporting information

S1 Table. Reaction number, descriptions, and BNGL code for the model reaction rules.

S2 Table. Species and initial concentrations.

S3 Table. Model parameters.

S4 Table. Nominal and bounds values for parameters in the PARPi Logic Gate Model.

S5 Table. Genes over- and under-expressed in ovarian spheroids treated with olaparib.

S1 Text. Model code.

S2 Text. Detailed discussion regarding upper and lower bounds for parameters utilized in the global sensitivity (Sobol) analysis.

S1 Fig. Three stage simulation of the Bogdal model: Replication of Fig 9B.

S2 Fig. Pfam domain structure of GSPT2.

S3 Fig. Growth inhibition of HCT116 colon tumor lines (WT or p53 null) exposed to selected clinical PARP1 inhibitors.

## Acknowledgements

SDM would like to thank Dr. Amanda Paulson for her efforts in coordinating laboratory experiments and contribution to review of the manuscript. We would also recognize Ms. Naomi Ohashi and Ms. Rebecca Lein for their project management roles at the Frederick National Laboratory for Cancer Research and ATOM Research Alliance.

## Author contributions

Conceptualization (overarching goals) – SDM, EAS

Data Curation (all aspects: creating, analyzing) – SDM, MPJ, MED

Formal Analysis (statistics, etc.) – SDM, NMI, BJY, KRR, NU

Funding Acquisition - EAS

Investigation (performing experiments; conducting research) – SDM, NMI, MPJ, BJY, NU, MRW

Methodology (creation of models) – SDM, EAS

Project Administration (overall planning and coordination) – SDM

Resources

Software

Supervision (oversight and leadership) – SDM, MED, NU, EAS

Validation (of findings, reproducibility) - SDM

Visualization (of the work) – SDM, NMI

Writing – Original Draft Preparation – SDM, NMI, MPJ

Writing – Review and Editing – SDM, NMI, MPJ, MED, BJY, KRR, NU, MRW, EAS

## Funding clause

This work represents a multi-institutional effort. Funding sources include the following: Lawrence Livermore National Laboratory internal funds; the National Nuclear Security Administration; and federal funds from the National Cancer Institute, National Institutes of Health, and the Department of Health and Human Services, Leidos Biomedical Research Contract No. 75N91019D00024, Task Order 75N91019F00134 through the Accelerating Therapeutics for Opportunities in Medicine (ATOM) Consortium under CRADA TC02349. This work was performed under the auspices of the U.S. Department of Energy by Lawrence Livermore National Laboratory [Contract No. DE-AC52-07NA27344]. (SDM, MPJ, MED, MRW, EAS)

This work was also supported by the New York Empire State Development Big Data Science Capital Project, the DOE Advanced Scientific Computing Research Applied Mathematics Fellowship, and the Brookhaven Laboratory Directed Research and Development DEDUCE Project. (NMI, KRR, BJY, NU)

## Competing interests

SDM is Founder and CEO of BioSystems Strategies, LLC.

## Notes

### Competing Interest Statement

Susan D. Mertins is Founder and CEO of BioSystems Strategies, LLC. All remaining authors report no competing interests.

