## Supplementary material for "Pharmacodynamic model of PARP1 inhibition and global sensitivity analyses can lead to cancer biomarker discovery": S1 Text

```
#
#
#                        This BioNetGen model code is described in the
article:
#
# "Pharmacodynamic Model of PARP1 Inhibition and Global Sensitivity
Analyses can Lead to Biomarker Discovery"
#           by  Mertins SD, Isenberg N, Reyes K-R, Urban N, Yoon B-J,
Jogalekar M, Diolaiti M, Ashworth A,
#                                       Weil MR, Stahlberg EA
#                                 (PLOS Computational Biology XXXX).
#
# In order to execute this model, please read the accompanying Model
Document file
# and consult the published article for a detailed description and
analysis.
#
#                                        ______INTRO________
#
#     This model follows the 11 ODE equations given in Bodgal, M et al.
2013 BMC Systems Biology
# that provides a minimal induction of apoptosis by p53 and AKT.
These two pathways can be
# activating or opposing apoptosis signaling.  It is notable that the
output is dependendent on p53, AKT,
# BAD, and BCLXL levels.  In the report, it is shown that at high
protein expression levels for BAD,
# an OR logic gate exists such that p53 OR AKT signaling can promote
apoptosis.  However, at
# lower protein expression levels for BAD, an AND gate exists such
that both signals are needed
# to trigger apoptosis.  The reaction rules below provide for the AND
logic gate.
#
#     In this model, unphosphorylated AKT is present as representing
growth factor withdrawl and therefore,
# primed apoptosis.  If sufficient p53 is present and bound to DNA
double strand breaks, caspase activation
# will occur.  Further, p53 and PARP1 are competitive for DNA double
strand breaks, and absent any
# PARP1 inhibitor (PARP1i), little or no apoptotic mediators are
present.  When PARP1i are bound,
# PARP1 is no longer available to bind DNA double strand breaks, there
is increased p53-DNA species and
```

```
# BAX is transcribed at a higher rate and triggers downstream caspase
activation.
#
#  The goal of the model was to input PARP1 and its inhibitor and to
output a therapeutic value. In this
# instance, the level of active caspase at the final steps or the
level of BAX can be considered therapeutic
# values.
#
# Note:  Well mixed reactions are considered and no cell volume is
inputted, but is taken into account
#
#                                   __________Model
Development____________
#
# Created in RuleBender 2.3.0
# BioNetGen 2.4.0
# MAC OS 10.15.7
# Susan D. Mertins, Ph.D.
# February 17, 2022
#
#                                   ____________BioNetGen
Model_____________
#
begin model

begin parameters

s1   1e-2  #Basal BaxmRNA synthesis rate
s2   3e-2  #p53 regulated BaxmRNA synthesis rate due to activation by
DNA Damage
s3   2e1   #Pro-caspase synthesis rate
s4   2e-1  #BAX protein synthesis rate

d1   1e-3  #1/s; BaxmRNA degradation rate
d2   1e-4  #1/s: BAX protein degradation rate
d3   2e-4  #1/s: Pro-caspase and caspase degradation rate

M    1.0e5  #Michaelis Menten Constant for p53 transcription rate

b1   3e-5  #molecules-1 sec-1:  BAX-BCLXL on rate
b2   3e-3  #molecules-1 sec-1:  BCLXL-BAD on rate
b3   3e-3  #molecules-1 sec-1:  phosphoBAD-Fourteen-three-three on
rate
```

```
b4   3e-5  #molecules-1 sec-1:  p53-DNA PARP-DNA on rate

u1   1e-4  #1/s:  BAX-BCLXL off rate
u2   1e-4  #1/s:  BCLXL-BAD off rate
u3   1e-4  #1/s:  phosphoBAD-Fourteen-three-three off rate
u4   1e-4  #1/s:  p53-DNA and PARP-DNA off rate

p1   3e-10 #1/s:  Kinase activity of AKT for BAD
q1   3e-5  #1/s:  Dephosphorylation rate for phosphoBAD
a1   2e-10 #molecules-1 sec-1:  Procaspase activation rate by BAX
a2   1e-12 #molecules-1 sec-1:  Procaspase autoactiavtion rate

kf1  1e-3  #molecules-1 sec-1: on rate for NAD binding
kr1  3.79  #1/s:  off rate for NAD based on reported Kd

IC50 0.001    #in uM; IC50 will vary as an input
kf2  1e-3     #in competition with NAD, therefore adjusted as above
kr2  IC50*3.79

kcat1  1   # Catalytic activity for PARylation is reported
(BindingDB.org)
kcat2  1   # Catalytic activity for de-PARylation is reported
(BindingDB.org)

DNADSBtot 1.74e5  #Adjusted for this model
AKTtot    0       #Value represents amount of active (phosphorylated)
AKT
p53tot    8.5e4   #This value is critical to demonstrated competition
with PARP-DNA binding
BADtot    0.6e5   #This level of BAD is found in the AND logic gate
(Bogdal, BMC Sys Biol 2013)
BCLXLtot  1e5     #This level of BCLXL is required for AND logic gate
(Bogdal, BMC Sys Biol 2013)
SCAFtot   2e5     #Found in Lipniaki PLoS Comp Biol 2016, SCAF =
Fourteen-three-three
PARPtot   1.07e5  #Estimage PARP concentration (Geiger et al.)
NADtot    1.07e6  #estimated on PARP concentration (i.e. one log
higher than PARP)
XRCC1tot  1.07e5  #estimated on PARP concentration
PARGtot   1.07e4  #relative to PARP as defined in vivo
Inhtot    0.52e5  #estimated as 0.5 of PARPtot

end parameters
```

```
begin molecule types

DNADSB(br)
p53(dna)
mRNA_Bax()
Bax(b)
BclXL(b)
Bad(S75_S99~0~PP,b)
Fourteen_3_3(b)
Caspase(csp~Pro~Act)
PARP(DNA,CD~inact~act,NAD)
NAD(sub)
XRCC1(Glu~uPAR~PAR)
PARG(CD)
Inh(isub)

end molecule types

begin seed species

DNADSB(br)                 DNADSBtot
p53(dna)                   p53tot
BclXL(b)                   BCLXLtot
Bad(S75_S99~0,b)           BADtot
Fourteen_3_3(b)            SCAFtot
PARP(DNA,CD~inact,NAD)     PARPtot
NAD(sub)                   NADtot
XRCC1(Glu~uPAR)            XRCC1tot
PARG(CD)                   PARGtot
Inh(isub)                  Inhtot

end seed species

begin observables

Molecules mRNA_Bax          mRNA_Bax()
Molecules Bax_free          Bax(b)
Molecules Caspase_act       Caspase(csp~Act)
Molecules p53_DNAbound      p53(dna!+)
Molecules PARP_Inhbound     PARP(DNA,CD~act,NAD!4).Inh(isub!4)

end observables

begin reaction rules
```

```
  # p53 binds DNADSB as representation of transcription factor
activity
  p53(dna) + DNADSB(br) <-> p53(dna!1).DNADSB(br!1) b4, u4

  # PARP binds DNADSB per function
  PARP(DNA,CD~inact,NAD) + DNADSB(br) <-> PARP(DNA!
2,CD~act,NAD).DNADSB(br!2) b4, u4

  # PARP binds substrate NAD
  PARP(DNA!2,CD~act,NAD).DNADSB(br!2) + NAD(sub) <-> \
  PARP(DNA!2,CD~act,NAD!3).DNADSB(br!2).NAD(sub!3) kf1, kr1

  # PARP PARylates XRCC1 as a representative substrate
  PARP(DNA!2,CD~act,NAD!3).DNADSB(br!2).NAD(sub!3) + XRCC1(Glu~uPAR)
-> \
  PARP(DNA!2,CD~act,NAD).DNADSB(br!2) + NAD(sub) + XRCC1(Glu~PAR)
kcat1

  # PARG de-PARylates XRCC1
  PARG(CD) + XRCC1(Glu~PAR) -> PARG(CD) + XRCC1(Glu~uPAR) kcat2

  # Inhibitor binds to NAD pocket and PARP falls off DNA.
  Inh(isub) + PARP(DNA!2,CD~act,NAD).DNADSB(br!2) <-> \
  Inh(isub!4).PARP(DNA,CD~act,NAD!4) + DNADSB(br) kf2,kr2

  # Baseline transcription and degradation of BAX mRNA
  0 <-> mRNA_Bax() s1, d1

  # p53 regulated BAX transcription via Hill function, typical rule
for transcription
  0 -> mRNA_Bax() s2*(p53_DNAbound^2/(M^2 + p53_DNAbound^2))

  # Bax translation, protein degradation
  0 <-> Bax(b)  s4*mRNA_Bax, d2

  # Bax--BclXL binding, unbinding
  Bax(b) + BclXL(b) <-> Bax(b!1).BclXL(b!1)  b1, u1

  # Bax (complexed) degradation
  Bax(b!1).BclXL(b!1) -> BclXL(b)  d2

  # BclXL and dep'ylated Bad binding, unbinding
  BclXL(b) + Bad(S75_S99~0,b) <-> BclXL(b!2).Bad(S75_S99~0,b!2)  b2,
```

```
u2

  # Bad p'ylation by AKT, dep'ylation
  Bad(S75_S99~0,b) <->  Bad(S75_S99~PP,b)  p1*AKTtot, q1

  # Bad (p'ylated) and 14-3-3 binding, unbinding
  Bad(S75_S99~PP,b) + Fourteen_3_3(b) <-> Bad(S75_S99~PP,b!
3).Fourteen_3_3(b!3)  b3, u3

  # BclXL unbinding from Bad upon Bad p'ylation by AKT.
  BclXL(b!2).Bad(S75_S99~0,b!2) -> BclXL(b) + Bad(S75_S99~PP,b)
p1*AKTtot

  # Unbinding of Bad from 14-3-3 upon Bad dep'ylation
  Bad(S75_S99~PP,b!3).Fourteen_3_3(b!3) -> Bad(S75_S99~0,b) +
Fourteen_3_3(b)  q1

  # Procaspase synthesis
  0 -> Caspase(csp~Pro)  s3

  # Caspase and procaspase degradation
  Caspase() -> 0  d3

  # Caspase activation by Bax and by other caspases
  Caspase(csp~Pro) -> Caspase(csp~Act)  a1*Bax_free + a2*Caspase_act^2

end reaction rules

end model

begin actions

generate_network({overwrite=>1})
simulate_ode({t_end=>600000,n_steps=>600})

end actions
```
