## Supplementary material for "Pharmacodynamic model of PARP1 inhibition and global sensitivity analyses can lead to cancer biomarker discovery": S2 Text

**S2 Text. Detailed Discussion regarding Upper and Lower Bounds for Parameters Utilized in the Global Sensitivity (Sobol) Analysis (See also S4 Table).**

**Parameter s1 for mRNA Synthesis Rate (Transcription)**

Measured values of mRNA elongation by RNA polymerases demonstrated that for eukaryotes, 20 - 100 nucleotides/s are transcribed. If the average gene length is $10^4$-$10^6$ nucleotides (nt) in eukaryotes (including introns), then calculations show that 1.5 min are needed to transcribe the $10^4$ nt gene. This translates to 0.6 molecules per min and this, in turn, equals 1e-2 mol/sec = s1.

It should be noted that 1e-2 mol/sec is expected to be the lower limit here but there is a multiplicative aspect of 5x given the measured range above of 20 - 100 nt/s. In sum, the upper and lower bounds for Parameter s1 is 1e-2 to 5e-2 mol/s (1–3).

**Parameter s2 for upregulated mRNA Synthesis Rate (Transcription)**

For Parameter s2, the mRNA synthesis rate is higher since it is upregulated by p53. But for this model, the nominal value is 3e-2 mol/s, then applying the same calculation from above, the lower and upper bounds would be 3e-2 to 1.5e-1 mol/s (1–3).

**Parameters s3 and s4 Protein Synthesis Rate (Translation)**

At BioNumbers.org, it was found that the protein synthesis rate for eukaryotes is approximately 2.8 - 10 aa/sec (2). Therefore, the length of the protein affects the synthesis rate.

For s3, pro-caspase, assumed to be caspase-3, is approximately 300 aa in length; therefore, it would require 30 – 107 sec to produce for one molecule or 0.009/s - 0.033/s. This range is significantly lower than the nominal value of 2e1 mol/s. Thus, the lower and upper bounds were established to be 9e-3 – 2e3/s. It would appear then that this is a log normal distribution.

For s4, BAX protein is approximately 192 aa; therefore, the range for one molecule synthesized is 19/s - 70/s, or 0.01/s - 0.05/s. This range is significantly lower than the nominal value of 2e-1 mol/s. Thus, the lower and upper bounds were established to be 1e-2 to 2e-1/s (2,4–6).

**Parameter d1 for BAX mRNA degradation rate**

An average lifetime for mRNA is between 3 and 8 min or 1 molecule degrades over 180 sec - 480 sec (2,7). Thus, the lower and upper bounds were established in the range of 2e-3 to 5e-3/s. Of note, the nominal value is 1e-3. These bounds are multiplicative with approximately a 5-fold multiplier.

**Parameter d2 and d3 for BAX, pro-caspase, and caspase protein degradation rates**

An average lifetime for a protein is 0.03 to 0.82 per hour (2,8). Proteins are expected to have longer lifetimes than mRNAs. Some proteins have exceptionally long lifetimes, but the average is useful for the model. Calculations show that the average lifetime translates to 8.3e-6/s - 2.16e-4/s. The nominal values (d2 and d3) for these proteins are 1e-4/s and 2e-4/s, respectively. Therefore, the lower and upper bounds are 8.3e-6 to 2.16e-4/s.

**Parameter M for Michaelis-Menten coefficient**

The Michaelis-Menten equation satisfactorily describes transcription mediated by factors and its activation coefficient ($K_m$) may take a wide range from 4e-1 μM - 8e3 μM (3,9). However, for this limited model the lower and upper bounds were established to be 9e4 to 1.1e5 μM.

**Parameters b2 and b3 for binding/on rate/association rate constant**

The nominal value for Parameters b2 and b3 is 3e-3/molecules/s. According to calculations from Sekar and Faeder (10), this binding rate, on rate, or association rate constant is considered fairly fast for protein-protein interactions. However, it is a critical aspect of the model such that BAD-BCLXL binding occurs more quickly when BAD increases in concentration to release BAX from BCLXL and trigger caspase activation. Therefore, the lower and upper bounds were established at 1e-3 to 1e-4/molecules/s.

**Parameter b1 for binding in protein-protein interactions, or on rate/association rate constant**

The nominal value for b1 is 3e-5/molecules/s. This is a rate constant is slower than the parameters b2 and b3 association rate constants above (10). As mentioned above, in the confines of the model, this is consistent. Therefore, the lower and upper bound were established at 1e-6 to 5.9e-5/molecules/s.

**Parameter b4 for binding in DNA/protein interactions, or on rate/association rate constant**

The nominal value (3e-5/molecules/s) for PARP1 and p53 binding DNA is the same because they both bind DNA. This is an important assumption since, in the model, PARP1 inhibition must activate caspase via p53 binding to DNA. Future models may account for the fact that p53 acts as a transcription factor and binds DNA at a distinct sequence and PARP1 binds to damaged DNA albeit *in vivo*, the DNA may have distinct sequence (for p53) and is nonspecifically damaged (for PARP1). The nominal value is again consistent with the modeling guidelines offered by Sekar and Faeder (10). Further, it is also important to mention that in a report by Lemersier and others, association rate constants for small molecule binding to a protein are approximately 1e-6/molecules/s (11). Therefore, the lower and upper bounds for Parameter b4 are 2.4e-5 to 3.6e-5/molecules/s.

**Parameters u1 - u3 for unbinding/off rate/dissociation rate constants in protein-protein interactions**

The nominal values are consistent with the association rate constants as described above for activation of caspase via p53-mediated transcription of BAX mRNA (10). Importantly, the lower and upper bounds for u1 is a larger range because BAX-BCLXL protein-protein dissociation is expected to occur at a faster rate (6e-5/s to 4e-4/s). For u2 and u3, the lower and upper bounds were 6e-5 to 1.4e-4/s.

**Parameter u4 for unbinding/off rate/dissociation rate constants in protein-DNA interactions**

The nominal value reflects weak interactions (u4) (10). It was chosen to maintain consistency within the model and to assume that when DNA damage occurs, PARP1 binds, repair occurs rather than act as a trigger for apoptosis mediated by p53. Importantly, PARP1 DNA binding domain affinity has been measured and is calculated from the Kd (12). Since the model assumes that p53 and PARP1 compete for DNA, u4 is utilized for both unbinding events and the lower and upper bounds were 6e-5 to 1.4e-4/s.

**Parameters p1 and q1 for catalytic rates of a kinase and phosphatase, respectively**

The nominal values are exceptionally slow but are consistent for the model. Measured turnover rates (or catalytic rates) for kinases and phosphatases can range from 0.057/s - 0.98/s as examples (See the BRENDA Database (13,14)). Therefore, the lower and upper bounds were established to have a normal distribution around the nominal values reflecting the mean (1e-11 to 9e-10/s for AKT kinase catalytic rate and 1e-6 to 9e-5/s for a generic dephosphorylation rate).

**Parameters a1 and a2 for pro-caspase and caspase activation rates**

This pair of parameters are likely to be a measurement of protease activity following a significant number of reactions that are not included in the model. The nominal values are exceptionally slow but are consistent. In the BRENDA Database, the measured turnover rates (of caspase activation or protease activity) range from 8.3e-3/molecules/s – 1.95e1/molecules/sec (13,14). For the purposes of establishing lower and upper bounds, a normal distribution around the nominal values was utilized (1.2e-11 to 2.8e-10/s for pro-caspase activation rate and 6e-13 to 1.4e-12/s for caspase activation rate).

**Parameters kf1 and kr1 for NAD binding and unbinding to PARP1 (on and off rates, association and dissociation rates)**

The kf1 association rate constant for NAD binding to xanthine dehydrogenase was measured and approximated the nominal rate of 1e-3/molecules/sec (15). Therefore, the lower and upper bounds were established as a normal distribution from 8e-4 to 1.2e-3/molecules/sec.

The kr1 dissociation rate constant was a measured value from benzamide binding to PARP1 (16). Similarly, the lower and upper bounds were considered a normal distribution and established as 3e0 to 4.58e0/s.

**Parameters kf2 and kr2 for generic Inhibitor binding and unbinding to PARP1 (on and off rates, association and dissociation rates)**

The kf2 association rate constant for Inhibitor binding to PARP1 was considered in competition with its substrate, NAD and thus equal to 1e-3/molecules/sec. As with NAD binding, the lower and upper bounds were considered a normal distribution and established to be 3e0 to 4.58e0/s.

In order to accommodate the inhibitory action of low dose IC50 values which demonstrate greater PARP1 inhibition, kr2 is a derived function dependent of kr1 (kr2 = IC50*3.79). The lower and upper bounds of the IC50 values were 1e-4 to 1e1 μM.

**Parameters kcat2 and kcat3 for catalytic rates (or enzymatic activity) of PARP1 and PARG**

The values set for the catalytic activity parameters resulted from previous parameter scans that indicated balanced rates were needed (1.0/s). Differential effects were accounted for in the initial concentrations of both enzymes. The BRENDA Database reports the measured ranges for PARP1 (2.0e-4/s - 1.2e1/s) and PARG (9.0e1/s - 2.6e2/s) (13,14). Therefore, the lower and upper bounds were considered a normal distribution and established at 6e-1 to 1.4e0/s.
