## Supplementary material for "Pharmacodynamic model of PARP1 inhibition and global sensitivity analyses can lead to cancer biomarker discovery": S1 Fig

### S1 Figure.

## A.

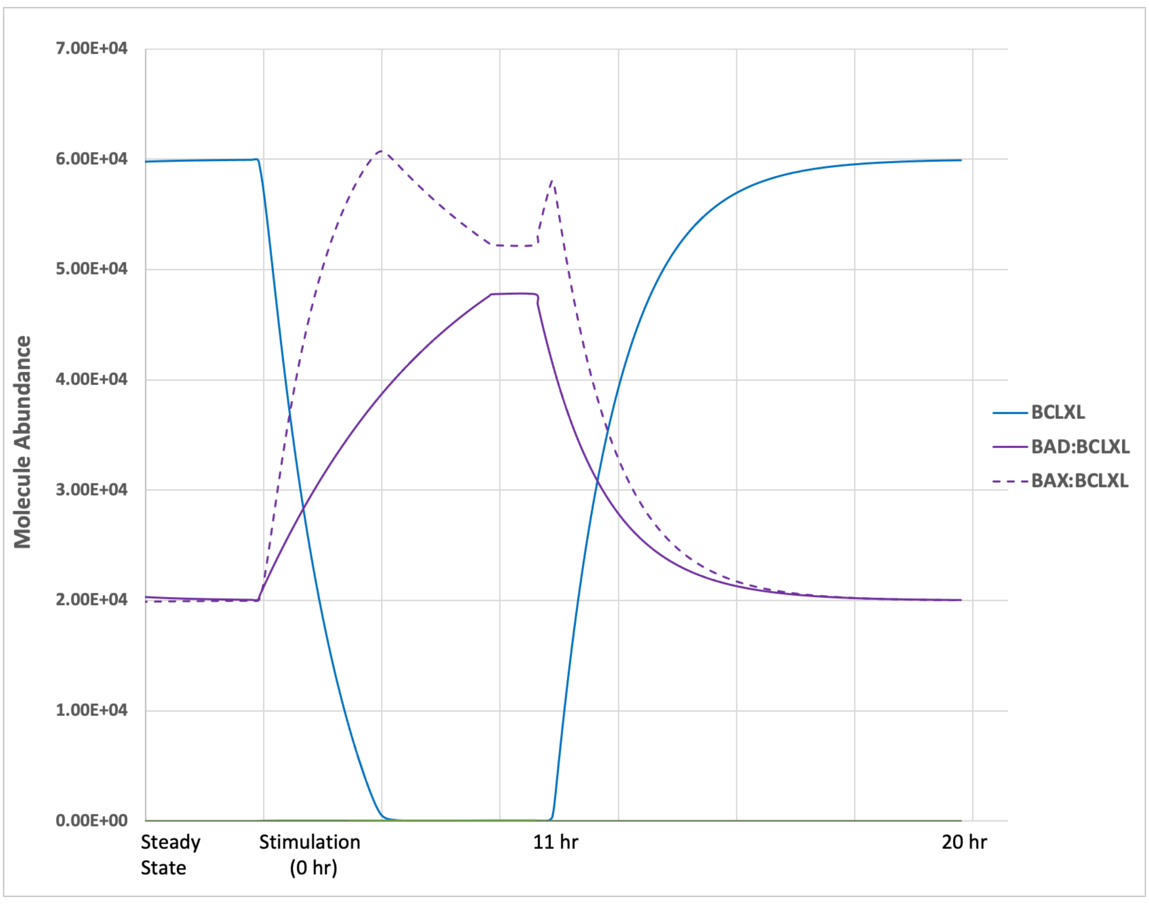


## B.

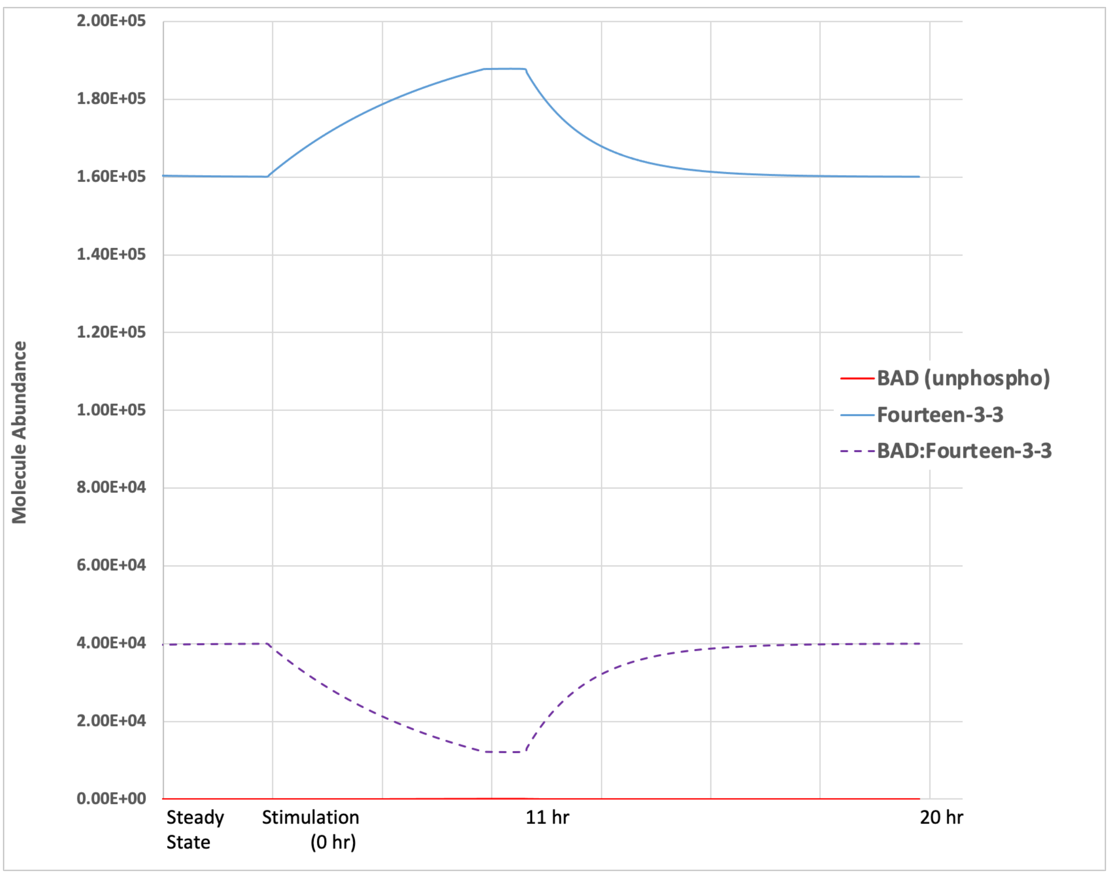


## C.

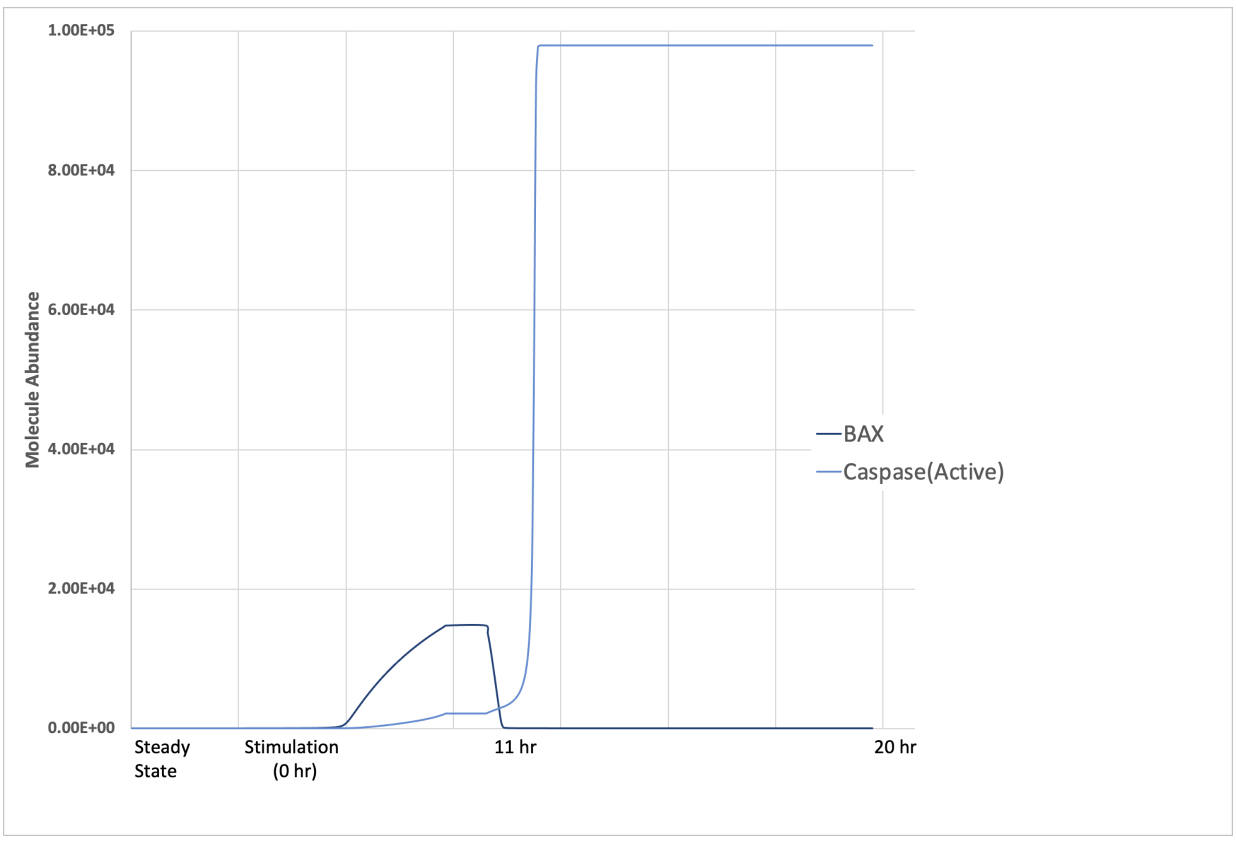


**S1 Figure. Three Stage Simulation of Bogdal et al. Model : Reproduction of Figure 9B.** The 11 ODEs in the report were coded in a rule-based model and simulated for 3 stages: steady state, induction of apoptosis (simulation) and relaxation. Figures S1A-C report molecule number for a given species listed in the legend. The stimulation phase included species concentrations for p53 and active AKT to induce caspase activation under the AND logic gate(C). All molecule abundances are identical for rule-based model and those in Figure 9B.
