## Supplementary material for "Pharmacodynamic model of PARP1 inhibition and global sensitivity analyses can lead to cancer biomarker discovery": S2 Fig

S2 Figure.

A.

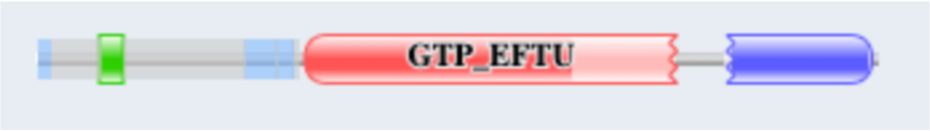


B.

| Source | Domain | Start | End |
|---|---|---|---|
| disorder | n/a | 1 | 52 |
| low_complexity | n/a | 2 | 10 |
| Pfam | PAM2 | 47 | 64 |
| disorder | n/a | 59 | 60 |
| disorder | n/a | 65 | 195 |
| low_complexity | n/a | 155 | 178 |
| low_complexity | n/a | 173 | 191 |
| Pfam | GTP_EFTU | 201 | 479 |
| disorder | n/a | 205 | 207 |
| disorder | n/a | 236 | 239 |
| disorder | n/a | 253 | 254 |
| disorder | n/a | 262 | 263 |
| disorder | n/a | 267 | 271 |
| low_complexity | n/a | 316 | 327 |
| Pfam | GTP_EFTU_D3 | 516 | 624 |

**S2 Figure. Pfam Domain Structure of GSPT2: Screenshots from Pfam Home (https://pfam.xfam.org).** Three domains in GSPT2 can be identified and 30/39 mutations described for TCGA uterine cancer patients fall within one of those domains. PAM2 domain binds PABP (HUGO ID HGNC: 8544), an important scaffold protein regulating GSPT2 function in both ribosome initiation and mRNA decay. GTP EFTU domain defines the P loop of the GTPase site. GTP EFTU D3 domain binds GSPT1 (or eRF3a, HUGO ID HGNC: 4621) thus regulating elongation termination.
