## Supplementary material for "Pharmacodynamic model of PARP1 inhibition and global sensitivity analyses can lead to cancer biomarker discovery": S3 Fig

S3 Figure.

a.

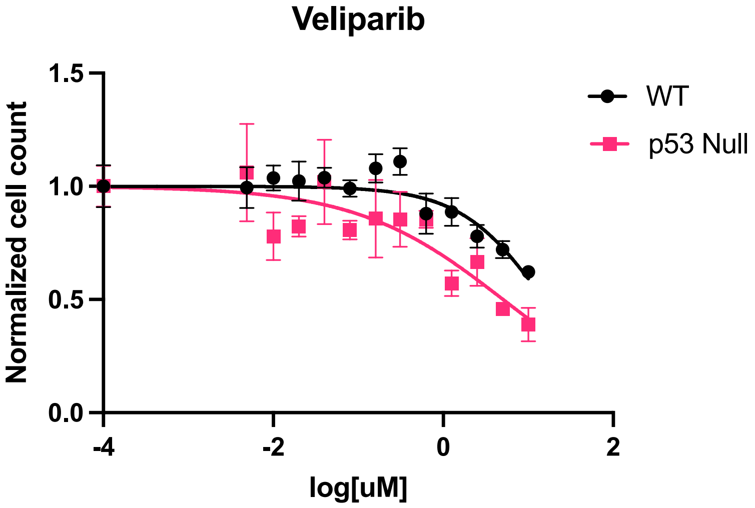


b.

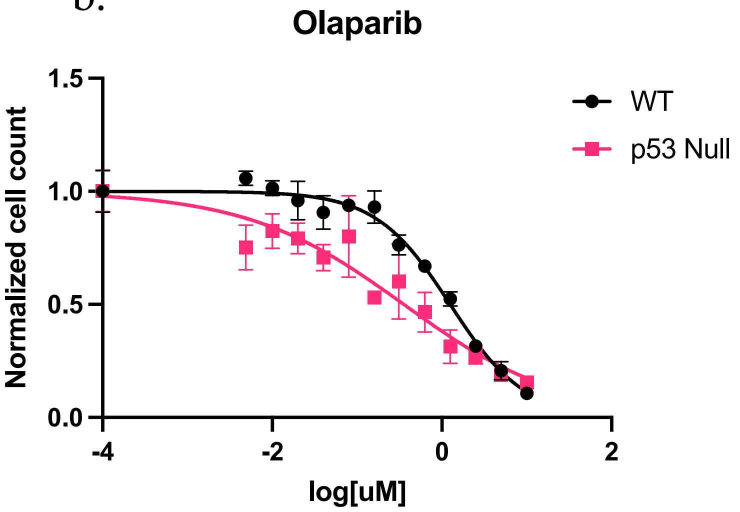


c.

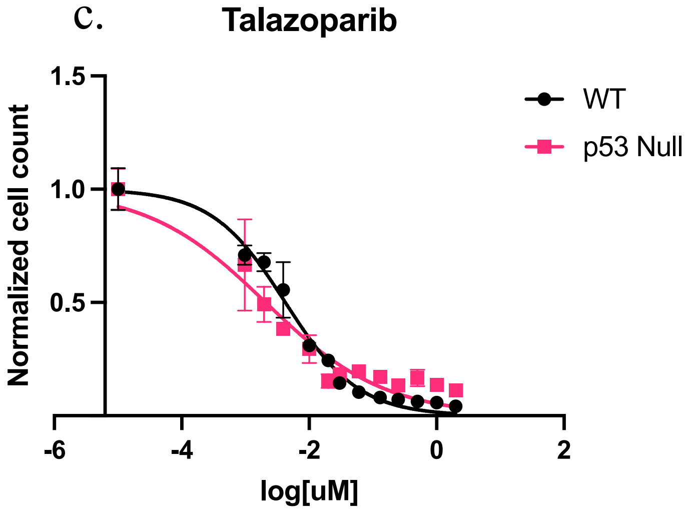


d.

| PARPi | WT | P53 -/- |
|---|---|---|
| Veliparib | > 10 uM | 5.07 uM |
| Olaparib | 1.25 uM | 0.355 uM |
| Talazoparib | 4.23 nM | 2.08 nM |

**S3 Figure. Growth Inhibition of HCT116 Colon Tumor Lines (WT or p53 null) Exposed to Selected Clinical PARP1 Inhibitors**. Survival curves of WT and p53 -/- cells treated with the indicated concentrations of (a) veliparib,(b) olaparib and (c) veliparib in a 92 hour assay Cell counts were measured using an incucyte zoom microscope and are plotted as mean values +/- SEM; n = 4. All values were normalized to DMSO control. (d) Table showing the concentration of each drug required to cause a 50% reduction in survival of WT or p53 -/- cells (SF50).
