## Supplementary material for "Pharmacodynamic model of PARP1 inhibition and global sensitivity analyses can lead to cancer biomarker discovery": S1 Table

**S1 Table. Reaction Number, Descriptions, and BNGL Code.**

| Reaction Number | Reaction Rule Description | BNGL Code |
|---|---|---|
| 1 | P53 binds double-stranded DNA break | `p53(dna) + DNADSB(br) <-> p53(dna!1).DNADSB(br!1)` |
| 2 | PARP1 binds double-stranded DNA break in the absence of NAD | `PARP(DNA,CD~inact,NAD) + DNADSB(br) <-> PARP(DNA!2,CD~act,NAD).DNADSB(br!2)` |
| 3 | PARP1 binds NAD | `PARP(DNA!2,CD~act,NAD).DNADSB(br!2) + NAD(sub) <-> PARP(DNA!2,CD~act,NAD!3).DNADSB(br!2).NAD(sub!3)` |
| 4 | PARP1 PARylates XRCC1 | `PARP(DNA!2,CD~act,NAD!3).DNADSB(br!2).NAD(sub!3) + XRCC1(Glu~uPAR) -> PARP(DNA!2,CD~act,NAD).DNADSB(br!2) + NAD(sub) + XRCC1(Glu~PAR)` |
| 5 | PARG dePARylates XRCC1 | `PARG(CD) + XRCC1(Glu~PAR) -> PARG(CD) + XRCC1(Glu~uPAR)` |
| 6 | Inhibitor binds to NAD pocket and PARP1 leaves DNA | `Inh(isub) + PARP(DNA!2,CD~act,NAD).DNADSB(br!2) <-> Inh(isub!4).PARP(DNA,CD~act,NAD!4) + DNADSB(br)` |
| 7 | Baseline transcription and degradation of BAX mRNA | `0 <-> mRNA_Bax()` |
| 8 | P53 mediated BAX mRNA transcription | `0 -> mRNA_Bax()` |
| 9 | BAX translation and protein degradation | `0 <-> Bax(b)` |
| 10 | BAX – BCLXL binding and unbinding | `Bax(b) + BclXL(b) <-> Bax(b!1).BclXL(b!1)` |

| | | |
|---|---|---|
| 11 | BAX – BCLXL complex degradation | `Bax(b!1).BclXL(b!1) -> BclXL(b)` |
| 12 | depBAD – BCLXL binding and unbinding | `BclXL(b) + Bad(S75_S99~0,b) <-> BclXL(b!2).Bad(S75_S99~0,b!2)` |
| 13 | BAD phosphorylation and dephosphorylation | `Bad(S75_S99~0,b) <-> Bad(S75_S99~PP,b)` |
| 14 | pBAD – Fourteen-3-3 binding and unbinding (dependent on mass action rates) | `Bad(S75_S99~PP,b) + Fourteen_3_3(b) <-> Bad(S75_S99~PP,b!3).Fourteen_3_3(b!3)` |
| 15 | pBAD – BCLXL unbinding | `BclXL(b!2).Bad(S75_S99~0,b!2) -> BclXL(b) + Bad(S75_S99~PP,b)` |
| 16 | pBAD – Fourteen-3-3 unbinding (dependent on dephosphorylation rate) | `Bad(S75_S99~PP,b!3).Fourteen_3_3(b!3) -> Bad(S75_S99~0,b) + Fourteen_3_3(b)` |
| 17 | Pro-caspase synthesis | `0 -> Caspase(csp~Pro)` |
| 18 | Caspase and pro-caspase degradation | `Caspase() -> 0` |
| 19 | Caspase activation | `Caspase(csp~Pro) -> Caspase(csp~Act)` |
