## Supplementary material for "Pharmacodynamic model of PARP1 inhibition and global sensitivity analyses can lead to cancer biomarker discovery": S2 Table

**S2 Table. Species and Initial Concentrations.**

| Species in BNGL Code | Annotation | Initial Concentration (molecules) |
|---|---|---|
| `DNADSB(br)` | DNA with break (`br`) as binding component | 1.74E5 |
| `p53(dna)` | P53 with component for DNA (`dna`) binding | 8.50E4 |
| `mRNA_Bax()` | BAX mRNA | Ø |
| `Bax(b)` | BAX with component for binding to BCLXL (`b`) | Ø |
| `BclXL(b)` | BCLXL with component for binding to BAX or BAD (`b`) | 1.00E5 |
| `Bad(S75_S99~0~PP,b)` | BAD with 2 components, amino acids S75 and S99 that have two states, phosphorylated and unphosphorylated (`S75_S99~Ø~PP`) and/or binding to BCLXL or Fourteen-3-3 (`b`) | 0.60E5 |
| `Fourteen_3_3(b)` | Scaffold protein, 14-3-3 with component for binding BAX (`b`) | 2.00E5 |
| `AKTtot` | AKT, a kinase | Ø for the AND Logic Gate. |
| `Caspase(csp~Pro~Act)` | Caspase with one component (`csp`) that can be in the inactive (`Pro`) or active (`Act`) state | Ø |
| `PARP(DNA,CD~inact~act,NAD)` | PARP1 with 3 components, one binds DNA (`DNA`), one is the active site (`CD`), and one for substrate or inhibitor binding (`NAD`). The active site has two states inactive (`inact`) or active (`act`). | 1.07E5 |

| | | |
|---|---|---|
| XRCC1(Glu~uPAR~PAR) | XRCC1 has one component (Glu), a glutamine that can be PARylated (PAR) or not (uPAR). | 1.07E5 |
| PARG(CD) | PARG has one component representing its active site (CD). | 1.07E4 |
| Inh(isub) | This is a generic inhibitor that binds to PARP1 component, NAD (isub) | 0.52E5 |
