## Supplementary material for "Pharmacodynamic model of PARP1 inhibition and global sensitivity analyses can lead to cancer biomarker discovery": S3 Table

**S3 Table. Model Parameters.**

| Parameter Symbol | Description | Value | Unit |
|---|---|---|---|
| s1 | Basal BAX mRNA synthesis rate | 1.0E-2 | molecules/sec |
| s2 | P53 mediated BAX mRNA synthesis rate | 3.0E-2 | molecules/sec |
| s3 | Procaspase synthesis rate | 2.0E1 | molecules/sec |
| s4 | BAX protein synthesis rate | 2.0E-1 | molecules/sec |
| d1 | BAX mRNA degradation rate | 1.0E-3 | $sec^{-1}$ |
| d2 | BAX protein degradation rate | 1.0E-4 | $sec^{-1}$ |
| d3 | Procaspase and caspase degradation rate | 2.0E-4 | $sec^{-1}$ |
| M | Michaelis Menten coefficient for p53-DNA bound regulated BAX transcription | 1.0E5 | molecules |
| b1 | BAX - BCLXL binding rate | 3.0E-5 | $molecules^{-1}$ $sec^{-1}$ |
| b2 | BAD - BCLXL binding rate | 3.0E-3 | $molecules^{-1}$ $sec^{-1}$ |
| b3 | pBAD - Fourteen_3_3 binding rate | 3.0E-3 | $molecules^{-1}$ $sec^{-1}$ |
| b4 | P53 – DNA binding rate and PARP1 – DNA binding rate | 3.0E-5 | $molecules^{-1}$ $sec^{-1}$ |
| u1 | BAX – BCLXL unbinding rate | 1.0E-4 | $sec^{-1}$ |
| u2 | BAD – BCLXL unbinding rate | 1.0E-4 | $sec^{-1}$ |
| u3 | pBAD – Fourteen_3_3 unbinding rate | 1.0E-4 | $sec^{-1}$ |
| u4 | P53 – DNA and PARP1 – DNA unbinding rate | 1.0E-4 | $sec^{-1}$ |
| p1 | Phosphorylation rate of AKT | 3.0E-10 | $sec^{-1}$ |
| q1 | Dephosphorylation rate | 3.0E-5 | $sec^{-1}$ |
| a1 | Procaspase activation rate by BAX | 2.0E-10 | $molecules^{-1}$ $sec^{-1}$ |
| a2 | Procaspase autoactivation rate | 1.0E-12 | $molecules^{-2}$ $sec^{-1}$ |
| kf1 | PARP1 - NAD binding rate | 1.0E-3 | $molecules^{-1}$ $sec^{-1}$ |
| kr1 | PARP1 - NAD unbinding rate | 3.79 | $sec^{-1}$ |
| IC50 | Half-maximal inhibitory constant for PARP1 enzymatic activity | 0.0001 – 10.0 | μM |
| kf2 | Drug – PARP1 binding rate | 1.0E-3 | $molecules^{-1}$ $sec^{-1}$ |
| kr2 | Drug – PARP1 unbind rate | IC50*3.79 | $sec^{-1}$ |
| kcat1 | PARP1 PARylation rate | 1.0 | $sec^{-1}$ |
| kcat2 | PARG dePARylation rate | 1.0 | $sec^{-1}$ |
