## Supplementary material for "Pharmacodynamic model of PARP1 inhibition and global sensitivity analyses can lead to cancer biomarker discovery": S4 Table

**S4 Table. Nominal and bounds values for parameters in the PARP1i Logic Gate Model.**

| Parameter | Description | Nominal Value | Lower Bound | Upper Bound |
|---|---|---|---|---|
| s1 | Basal BAX mRNA synthesis rate | 1e-2 mol/s | 1e-2 | 5e-2 |
| s2 | p53 regulated BAX mRNA synthesis rate | 3e-2 mol/s | 3e-2 | 1.5e-1 |
| s3 | Pro-caspase protein synthesis rate | 2e1 mol/s | 9e-3 | 2e1 |
| s4 | BAX protein synthesis rate | 2e-1 mol/s | 1e-2 | 2e-1 |
| d1 | BAX mRNA degradation rate | 1e-3 $s^{-1}$ | 1e-3 | 5e-3 |
| d2 | BAX protein degradation rate | 1e-4 $s^{-1}$ | 8.3e-6 | 2.16e-4 |
| d3 | Pro-caspase and caspase protein degradation rate | 2e-4 $s^{-1}$ | 8.3e-6 | 2.16e-4 |
| M | Michaelis-Menten coefficient for p53 - DNA bound regulated BAX transcription | 1e5 | 9e4 | 1.1e5 |
| b1 | BAX - BCLXL binding rate | 3e-5 $mol^{-1}$ $s^{-1}$ | 1e-6 | 5.9e-5 |
| b2 | BAD - BCLXL binding rate | 3e-3 $mol^{-1}$ $s^{-1}$ | 1e-4 | 3e-3 |
| b3 | pBAD - Fourteen_3_3 binding rate | 3e-3 $mol^{-1}$ $s^{-1}$ | 1e-4 | 3e-3 |
| b4 | p53 - DNA and PARP1 - DNA binding rate | 3e-5 $mol^{-1}$ $s^{-1}$ | 2.4e-5 | 3.6e-5 |
| u1 | BAX - BCLXL unbinding rate | 1e-4 $s^{-1}$ | 6e-5 | 4e-4 |
| u2 | BAD - BCLXL unbinding rate | 1e-4 $s^{-1}$ | 6e-5 | 1.4e-4 |
| u3 | pBAD - Fourteeen_3_3 unbinding rate | 1e-4 $s^{-1}$ | 6e-5 | 1.4e-4 |
| u4 | p53 - DNA and PARP1 - DNA unbinding rate | 1e-4 $s^{-1}$ | 6e-5 | 1.4e-4 |
| p1 | Phosphorylation rate of AKT | 3e-10 $s^{-1}$ | 1e-11 | 9e-10 |
| q1 | Dephosphorylation rate | 3e-5 $s^{-1}$ | 1e-6 | 9e-5 |
| a1 | Pro-caspase activation rate by BAX | 2e-10 $mol^{-1}$ $s^{-1}$ | 1.2e-11 | 2.8e-10 |
| a2 | Pro-caspase autoactivation rate | 1e-12 $mol^{-1}$ $s^{-1}$ | 6e-13 | 1.4e-12 |
| kf1 | PARP1 - NAD binding rate | 1e-3 $mol^{-1}$ $s^{-1}$ | 8e-4 | 1.2e-3 |
| kr1 | PARP1 - NAD unbinding rate | 3.79 $s^{-1}$ | 3.0 | 4.58 |
| kf2 | PARP1 - Inhibitor binding rate | 1e-3 $mol^{-1}$ $s^{-1}$ | 8e-4 | 1.2e-3 |
| kr2 | PARP1 - Inhibitor unbinding rate dependent on IC50 | IC50*3.79 | 3.79e-3 | 37.9 |
| kcat1 | PARP1 PARylation rate | 1.0 $s^{-1}$ | 6e-1 | 1.4 |
| kcat2 | PARG dePARylation rate | 1.0 $s^{-1}$ | 6e-1 | 1.4 |
