## Supplementary material for "Pharmacodynamic model of PARP1 inhibition and global sensitivity analyses can lead to cancer biomarker discovery": S5 Table

**S5 Table. Genes Over- and Under-expressed in Ovarian Spheroids Treated with Olaparib (Taken from Sheta et al., 2020).**

Over Expressed Genes

VIM
SPRY2
GPC6
GLT8D2
CNRIP1
BX337332
DOK6
THC2574606
PECI
ZFP82
ROBO1
C6orf159
GFPT2
CCDC74B
HMGA2
CDO1
BAG2
NDN
NLGN1
SGCG
IL11
ZSCAN18
TCF4
BC039411
ZFP82
NBEA
THC2655811
TXNDC13
AK094088
X15674
QPCT
GPR84
DDR2
ITGA1
FAM171B
SNHG5
ZNF71
DPYSL4
TIMP1
AMIGO2
TRPS1
SH3BGRL
POPDC3
RBPMS2
CMTM7
STAU2
RASGEF1A
FBN1
AL831999
GJA1
SACS
FANCF
TMSB15A
THC2536817
ZNF347
MGAT3
MLLT11
BC013389
GAMT
A_32_P131583
TXNDC13
BEX4
LPHN2
GNB4
DKFZp547K054
BQ272125
FERMT2
C12orf39
TEX28
HOXB2
SLC2A14
ANKRD45
LOC400657
C3orf18
ZNF415
ZNF667
CLGN
C1orf38
BG185346
PLEKHO1
AW268902
CN312045
EPB41L3
AK125077
SLIT2
THC2675966
DDR2
ZC4H2
CAND2
MEIS1
ADAMTS6
AXL
GSPT2
CPNE8
MYL9
FLRT2
A_24_P926993
LOC283658
FAM69B
PLCB1
TMEM121
EXOSC6
KCNJ12
ANXA6
CR615016
RGS18
SNAI2
LOC641298
SH2B3
LIX1L
SPRY4
ESPNL
AU158345
FGFR1
UAP1
CPT1C
GLI2
EID3
KYNU
CDC42EP2
HOXB3
LGALS1
MAGED4B
FLJ44253
THC2654608
GPR125
DNAJC1
PDE4A
REEP2
C16orf45
DYRK3
B4GALT6
THC2497772
ETV5
DZIP1
AK130366
CMTM3
FBXO2
ABL2
SUSD5
BC041913
ENST00000377006
IPW
ITGAV
C16orf48
LIX1L
JAM3
C1orf9
LEPREL2
ENST00000285206
DCP1B
HOXB4
RGMB
LOC387895
STEAP2
SPAG7
PNMA1
KLF6
SLC25A37
RPA4
CDK2AP1
RBM12
TTC28
CEP170
IFFO1
EMP3
LOC100131482
AW389821
PDLIM7
SHC1
B4GALT3
NEO1
IPO5
DYX1C1
MAP3K12
WBSCR27
PCDHA11
LMO4
THC2709260
THC2531204
HTR7
RAPGEF2
AF086071

C18orf19
TEKT4
DUSP22
FSD1
BMI1
PTDSS2
NAP1L1
ZNF214
GNAZ
A_24_P324424
PHTF1
YRDC
DYNC1I2
FJX1
CCNG1
SNAPC2
ANKS3
KCNG1
AK022030
RPS6KA6
FBXL7
U69195
THC2500271
ST3GAL2
MRAS
THC2608658
CR598364
PTP4A2
NUDT11
ATXN2
ZMYM4
THC2609092
PDLIM7
TTC7B
DBN1
PTPN12
RTCD1
BTN2A1
KLHDC10
DOCK7
SEC23A
PCTK2
LMAN1
WRB
DAXX
PAPSS1
TNFRSF10B
BMI1
DENND5A
ZNF773
A_24_P238377
KIF1B
FARP1
EIF4EBP1
CXXC1
ST3GAL3
MFAP2

Under Expressed Genes

OSBPL3
IER5
CCDC24
STAT4
ME1
ZNF219
APOL1
HIST1H2AD
HIST1H2AJ
USP15
STAP2
CAPN2
SH3D19
MUDENG
DIRC2
AIFM2
ENST00000399573
EXOC3
KIAA0513
HIST2H2AC
COPZ2
SLC39A11
PKP2
SECTM1
BQ717518
EFNB2
KLF7
STXBP5
LRRC8E
HIBADH
PGM2
STAT1
PHKB
GPC1
ADI1
C20orf24
ASAP2
RNF217
ARAP3
AVPI1
LMNA
HIST3H2A
SAV1
ACOT8
ATP9A
CD14
PIK3IP1
LOC440894
ENST00000398832
CBLB
ADAL
NPEPL1
SGMS2
PLEKHA6
THC2619531
LRRFIP1
THC2596442
TNFRSF10A
IFITM2
ITM2B
PSG7
MEGF6
CTNNAL1
ATP5E
NMNAT3
CAMK2D
TAP1
ANKRD35
ANXA2P3
SC5DL
A_24_P868905
DUSP14
PLEKHH2
CAT
A_32_P167592
FAM110C
PLEKHA1
FGF18
MAP2K3
JUB
A_24_P101742
RNF125
MORC4
SULT1A1
ABCB1
DDX60
HSPA2
EIF2AK2
THC2507792
TNS3
PANK3
PARP12
SULT1A2
HIST1H1E
STARD5
MOSPD2
ITGB8
HLA-DRB1
MAPK13
HYLS1
THC2543840
ARID1B
LGALS8
CAST
DBI
YPEL3
HLA-G
ANXA2P1
HLA-C
SYNGR2
ANXA2
CXorf39
GALNAC4S-6ST
RNF141
MLKL
CCNYL1
A_23_P170713
CD109
PPIC
HIST1H2AG
A_24_P106166
IFITM4P
A_24_P332953
FLJ31356
MPZL3
OSBPL10
LOC646609
HIST1H2AH
SLC9A3R1
CAPG
SETD6
OXTR
A_32_P327750
THC2780391
SLC2A5
MYOF
FLJ21075
LOC341230
MAST4
CTSB

| | | | | |
|---|---|---|---|---|
| CR624517 | DNAJA4 | A_24_P358606 | BST2 | A_24_P890995 |
| KTN1 | CCDC125 | B4GALT1 | MATN2 | F11R |
| HIST1H2AM | GSC | IFITM3 | CARD10 | DDR1 |
| OPN3 | ATP1B3 | KRT8P15 | FAM25C | GALNT14 |
| TBC1D2 | ENST00000270031 | SLC48A1 | AK026984 | A_23_P125109 |
| PSG1 | PBX1 | IL1RAP | EGFLAM | MAF |
| CAMK2N1 | OSTF1 | TMEM187 | PYDC1 | PYCARD |
| AP1S3 | ULBP2 | TLR3 | EPS8L2 | NCOA7 |
| MYOF | SLC25A27 | S100A16 | APOL3 | S73202 |
| SASH1 | GPR126 | CAV2 | NAPRT1 | LTBR |
| THC2739760 | KRT23 | STK17A | A_24_P93111 | SAMHD1 |
| STAMBPL1 | A_24_P7040 | RNF43 | NP101106 | SEMA5A |
| PSD4 | SEMA3E | LAMC2 | ARHGEF16 | PLS3 |
| AHNAK | SSH3 | SMAD3 | INPP1 | SLC44A3 |
| TDRD7 | AK092921 | TRIM69 | HLA-F | GALNTL4 |
| HIST1H2AK | A_24_P900555 | LOC728323 | CARD17 | MET |
| P4HTM | NR1D2 | CD47 | CXADR | MYOM2 |
| C10orf65 | ARHGEF3 | CKB | ACP5 | THC2666453 |
| TST | CCL26 | BC063022 | BC006271 | ETV7 |
| GPRC5A | GSDMD | PAPSS2 | FLJ38359 | PLA2G16 |
| KIAA1147 | LAMC2 | LMO2 | OAS3 | JAZF1 |
| VAMP8 | ARHGEF5 | HLA-DRB3 | DDX58 | ZNF365 |
| NPC2 | KCTD12 | IFNGR1 | ARHGAP23 | CHADL |
| ARHGAP25 | WNT4 | LGALS8 | FBXO25 | RBM47 |
| SLC39A4 | AA455656 | hCG_1988300 | CBR3 | SNCA |
| F2RL1 | TRIM59 | EIF4E3 | SYNGR2 | A_24_P195974 |
| TPM1 | AW804939 | FLJ43692 | DNER | SAMD9L |
| GCOM1 | MAFB | KCNK2 | BIK | GLRB |
| TNFAIP2 | MITF | TOM1L2 | CMYA5 | TRIM38 |
| CR603450 | HLA-B | EY892390 | LCMT2 | PCSK1 |
| MR1 | KHDC1 | AF143325 | ITGB6 | SMARCA2 |
| PAG1 | SIPA1L2 | S100A11 | TREX1 | HSPB1 |
| ARNTL2 | CLIC3 | AL832534 | RIN2 | NUDT7 |
| CBR1 | C4orf32 | ANXA4 | NMI | DHRS3 |
| MALAT1 | LMO7 | HLA-C | NSUN7 | CR622110 |
| MGST2 | BCO2 | H2AFJ | ARSI | EZR |
| TMEM173 | TRIM14 | SKAP2 | THC2656519 | PDLIM1 |
| CD47 | ETS2 | C2orf55 | AK124281 | AHR |
| SP110 | A_24_P367100 | AK091178 | HIST1H1C | VAV1 |
| PTPN3 | THC2550342 | FRAS1 | AP1S3 | HERC6 |

| | | | | |
|---|---|---|---|---|
| ZNF594 | LIMCH1 | BC004287 | CASP1 | PSG8 |
| PVRL3 | AP1M2 | FAM134B | SULF2 | CPVL |
| MAP7 | NR3C1 | CYP39A1 | NFIA | A_24_P16230 |
| ATP1B1 | BTBD6 | GCNT1 | HTRA1 | hCG_1815491 |
| LOC100128918 | MYO5B | LOC149501 | PRR15 | TXNIP |
| CCDC144B | EPS8 | A_24_P15973 | A_24_P281374 | A_24_P6850 |
| A_24_P745352 | A_24_P178167 | PDE5A | RSAD2 | FLJ22662 |
| TMEM125 | PSG9 | SMPDL3A | C10orf116 | AI311458 |
| OTUD1 | EDN1 | A_32_P122492 | ADAL | IL1RAP |
| LOC644189 | MICALCL | GABRE | DEFB1 | LOC647954 |
| MAOA | B2M | PLEKHG4 | PLSCR1 | A_24_P109661 |
| MYLIP | PSG6 | SSFA2 | KLF5 | LAMA3 |
| HSPA1A | PSG4 | ARNT2 | KRT33A | ADM |
| THC2736113 | A_23_P370707 | AREG | HLA-F | KRT18P34 |
| CDK6 | CCDC144B | BC035647 | KRT18P40 | NTN4 |
| THC2659853 | C11orf52 | DSC3 | PRAGMIN | SQRDL |
| THC2697511 | ZSCAN16 | CCL28 | MX1 | KRT18P19 |
| ACOT4 | DENND2D | CTSH | AF390550 | GALNT3 |
| C9orf3 | AW444553 | AF088076 | PSMB8 | A_24_P409420 |
| CXADRP2 | THC2639718 | KCNK1 | IFIH1 | A_24_P230466 |
| F3 | THC2657193 | ZFP36L2 | A_24_P780319 | LOC442249 |
| ENST00000318291 | RASSF9 | CR614418 | KRT18P28 | THC2645298 |
| ANXA8L2 | LRRC61 | SAMD9 | MBP | CLDN1 |
| DOK7 | A_24_P418216 | SDPR | GLUL | A_24_P264293 |
| ARRDC4 | SCHIP1 | AF026246 | AA837799 | A_24_P383660 |
| MCTP2 | BC031342 | KIAA1324L | PTPN3 | A_24_P358406 |
| KRT8 | PARP9 | LIPH | DDX43 | CDKN2B |
| HCP5 | MARVELD2 | C10orf67 | KRT18P33 | P2RY2 |
| SLFN11 | C2orf74 | LOC26010 | LGALS3 | ARHGAP29 |
| FGD4 | C1orf21 | COL8A2 | AADAC | CDKN2A |
| PSMB9 | A_24_P281605 | C19orf33 | FAM129A | KRT18P30 |
| SNORD123 | CMAH | CD9 | A_24_P686014 | A_24_P24645 |
| SDR42E1 | LEPREL1 | A_32_P73413 | FAM84B | AK098314 |
| PLK2 | A_24_P401124 | A_32_P75141 | CTAGE4 | THC2524582 |
| NRP1 | C1orf226 | A_24_P384369 | A_24_P230486 | TMEM154 |
| TMPRSS3 | SGK1 | A_24_P264644 | PHLDB2 | PKIB |
| RHOD | PRSS23 | ITGA6 | CARD16 | NCF2 |
| IRF7 | THBD | ENST00000311208 | LAMB1 | A_24_P281443 |
| HLA-A | A_24_P341546 | DUSP23 | KRT18P49 | KRT18 |
| NAP5 | A_23_P72014 | C1orf59 | AK2P2 | EPB41L4A |

| | |
|---|---|
| IFI16 | JAG1 |
| LOC100128116 | PLAT |
| A_24_P161827 | CDH1 |
| MAL2 | |
| A_24_P418687 | |
| KRT18P42 | |
| A_24_P584463 | |
| ARMCX3 | |
| RTKN2 | |
| A_24_P306704 | |
| A_24_P471242 | |
| A_24_P161733 | |
| THC2535223 | |
| BMP4 | |
| A_24_P792988 | |
| IL18 | |
| FLJ40504 | |
| S100A10 | |
| TGM1 | |
| KRT19 | |
| PODXL | |
| BF803942 | |
| FST | |
| A_24_P247233 | |
| KCNS3 | |
| ANXA3 | |
| S100A6 | |
| TPD52L1 | |
| PERP | |
| A_32_P218707 | |
| SLPI | |
| LYPLAL1 | |
| KIAA1199 | |
| CD24 | |
| AK094799 | |
| GPX1 | |
| MTSS1 | |
| METTL7A | |
| ANXA1 | |
| EFEMP1 | |
